# Maternal prenatal stress induces sex-dependent changes in tRNA fragment families and cholinergic pathways in newborns

**DOI:** 10.1101/2024.07.10.602894

**Authors:** Shani Vaknine Treidel, Silvia M. Lobmaier, Ritika Sharma, Nimrod Madrer, Dana Shulman, Pnina Greenberg, Estelle R. Bennett, David S. Greenberg, Adi Turjeman, Camilla Zelgert, Peter Zimmermann, Martin G. Frasch, Liran Carmel, Marta C. Antonelli, Hermona Soreq

## Abstract

Maternal perceived prenatal stress (PPS) is a known risk factor for diverse developmental impairments in newborns, but the underlying molecular processes are incompletely understood. Here, we report that maternal PPS altered the birth profiles of blood transfer RNA fragments (tRFs), 16-50nt long non-random cleavage products of tRNAs, in a sex-dependent manner. Importantly, comparing stressed versus control maternal and umbilical cord blood serum presented alterations that were not limited to individual tRFs, but rather reflected selective changes in particular tRF families grouped by their mitochondrial or nuclear genome origin, parental tRNA coded amino acid, and cleavage type. tRF families that show stress- and sex-specific effects, revealed shared length and expression patterns which were strongest in the female newborns. Several of these tRFs carry complementary motifs to specific cholinergic mRNAs, suggesting possible translational regulation similar to microRNAs. Compatible with the cholinergic regulation of stress reactions, those “CholinotRFs” achieved an AUC of 95% when classifying female newborns according to maternal PPS. Moreover, we found altered catalytic activity of serum acetylcholinesterase, which was particularly elevated in male newborns, marking a second sex-specific effect. Our findings demonstrate an association of tRF families’ patterns with newborns’ sex-specific stress response to PPS and may lead to better diagnosis and therapeutic tools for these and other stressors.

## INTRODUCTION

The womb environment plays a crucial role in determining the future physical and mental health of the fetus, through the effects of numerous factors affecting fetal development (1–3). Specifically, psychosocial stress expressed as clinical or preclinical prenatal depression or anxiety affects 20% of all pregnancies (4–6). As psychological stress is difficult to measure directly due to its subjective nature, it can be evaluated using perceived stress measurement (6,7). Indeed, prenatal perceived stress (PPS) associated with offspring neurodevelopmental impairments, including neurobehavioral problems and greater vulnerability towards developing psychiatric disorders later in life (8,9). Moreover, PPS was negatively correlated with fetal left hippocampal volume (10,11), infant cognitive performance at 18 months (4), and newborn telomeres length (12,13). Some of these effects are sex-specific, pointing at male newborns as more vulnerable (5,14). Proposed mechanisms mediating these effects include epigenetic programing (15) and maternal immune activation through the hypothalamic-pituitary-adrenal axis (HPA), although cortisol-mediated fetal programming presented conflicting evidence (6,16,17). However, little is known about the role of short noncoding RNAs in these processes.

tRNA fragments (tRFs) are 16 to 50 nucleotides long noncoding RNAs which are highly conserved across the evolutionary tree and are cleaved from pre- and mature tRNAs by specific endonucleases (18,19). Human tRNA genes originate from both the nuclear and the mitochondrial (MT) genomes, and both are important for healthy cell function (20). Moreover, the nuclear genome contains “lookalike” tRNA genes, mimicking MT tRNAs, hailing their functional importance also outside the mitochondria (21). The more widely studied tRFs are cleaved from mature tRNAs, and are classified into five groups based on their cleavage point: 5’-half and 3’-half tRFs are produced by angiogenin cleavage at the anticodon loop, splitting the tRNA into two halves, whereas 5’-tRFs, i-tRFs, and 3’-tRFs are cleaved around the D- and T-loops by a combination of angiogenin, dicer and other endonucleases, splitting the tRNA into three parts (20,22). Interestingly, these cleavage points are not precise and can vary across tRFs sharing their cleavage type (20), sometime influenced by pre-existing tRNA modifications, such as methylation and pseudouridylation, which are actively involved in the biogenesis of tRFs (19,23). Further, tRFs can regulate transcription and translation by various mechanisms, including complementary binding to other RNAs or RNA binding proteins, promotion of ribosome biogenesis, and post-transcriptional gene silencing via the RISC complex (19,22–24). The roles of tRFs have been explored in neurodegenerative diseases, cancers, stroke and non-alcoholic fatty liver disease (23,25–28), but their impact on pregnancy complications or developmental disorders was only scarcely studied (29,30).

The cholinergic system may actively contribute to the above processes. It controls cognitive processes, inflammatory events, and neuromuscular communication (31–34). Further, it is a major regulator of stress responses, which play an active role in trauma reactions, neurodegeneration, stroke and other mental conditions (35,36). As a major player in both the central and the peripheral nervous system, acetylcholine is produced in neurons as well as in liver and immune cells and is hydrolyzed by two cholinesterase enzymes. The major cholinesterase in the brain is acetylcholinesterase (AChE) and in the periphery, butyrylcholinesterase (BChE) (31,32). Altered balance between the circulation activities of these two enzymes reflects different conditions, from anxiety to Alzheimer’s disease (35). Cholinesterases and other cholinergic genes are regulated in several psychological disorders by “CholinomiRs” (37), microRNAs (miRs) carrying complementary sequences to those of cholinergic genes that enable them to suppress their expression (32,38,39). Interestingly, tRFs may likewise regulate cholinergic genes via similar mechanisms. Importantly, circulating “CholinotRF” levels are elevated while CholinomiR levels decline in nucleated blood cells from post-ischemic stroke survivors (27), and mitochondrial-originated CholinotRFs show sex-specific cognition-related declines in the nucleus accumbens of women living with Alzheimer’s disease (25).

Based on these and other studies, we explored fetal reactions to PPS as part of the ‘FELICITy’ study collection, under the hypothesis that CholinotRFs may present a promising avenue to study newborns reactions to PPS (40–43). This unique cohort spanned a comprehensive battery of biomarkers in 120 mother-baby dyads, aiming to identify the stressed newborns as early a stage as possible. Previous studies yielded a list of biomarkers: the fetal stress index (FSI), measuring fetal heart rate reaction to fluctuations in the maternal heart rate during the third trimester; maternal hair cortisol levels at birth (43); umbilical cord serum (UCS) iron homeostasis markers (42); and DNA methylation in newborns’ saliva (41). All of these biomarkers presented differences between stress and control groups, some in a sex-specific manner, e.g., transferrin saturation levels which were changed only in male newborns in the PPS group (42).

Here, we describe an in-depth exploration of the impact of PPS on the newborn. To reflect maternal PPS assessed during the third trimester, we profiled modulated levels of tRFs and catalytically active cholinesterase enzymes in mothers and offspring at birth, as the earliest indicators of newborns’ reaction to PPS. These analyses identified sex-specific differences in particular tRF families as well as in cholinesterase activities, which differentiated between newborns of stressed and non-stressed mothers. Importantly, our analysis pipeline showed that grouping tRFs into families based on shared properties was more informative than considering each tRF type separately and that CholinotRFs enabled accurate PPS-based classification of female newborns.

## MATERIALS AND METHODS

### Participant recruitments and sample collection

A cohort of 18 - 45 years old women with singleton pregnancies was recruited (July 2016 - May 2018) by the Department of Obstetrics and Gynecology at Klinikum Rechts der Isar Hospital of the Technical University of Munich, as part of the FELICITy project (42,43). PPS was assessed during their early third trimester using the Cohen Perceived Stress Scale (PSS-10) questionnaire (7). Based on a pilot study, women who achieved PSS < 19 were assigned to the control group and those with PSS ≥ 19 to the PPS group (43). The women also completed the prenatal distress questionnaire (PDQ) (44) that assesses pregnancy-related stress and was highly correlated with PSS-10 scores (Supplementary Figure 1c). 728 women completed the PSS-10, and at the day of the birth venous maternal blood was collected from 128 of them upon entry into the delivery room. Umbilical cord blood (arterial or venous) was collected from 120 of the matched newborns post-partum. All blood samples were collected into 9 mL S-Monovette® Serum Gel with Clotting Activator tubes (Sarstedt, # 02.1388). Serum was separated by a 10 minutes centrifugation at 2500 RPM, frozen at -20°C and kept long-term at -80°C. Samples were shipped on dry ice to the Soreq lab at the Hebrew University of Jerusalem, where they were kept at -80°C until analysis.

### Cholinesterase activity measurements

Cholinesterase activity was measured using Ellman’s assay (45), in serum samples from 70 mothers and newborns who experienced non-emergency vaginal birth, to prevent birth types from confounding the results (58.3%, Supplementary Figure 1b). Briefly, each sample was defrosted rapidly in a water bath, diluted 1:20 in phosphate buffered saline (PBS), and 10 µL were mixed with 180 µL Ellman’s solution in 96 flat-well plates. Tests were all performed in triplicates. Each plate was incubated either without inhibitors, or with the BChE inhibitor tetraisopropylpyrophosphoramide (iso-OMPA; final concentration 50 µM; Sigma, T1505), or the AChE inhibitor BW284c51 (final concentration 5 µM; Sigma, A9103). To enable comparison between plates, each plate included four internal controls: (1) a random maternal sample, (2) its corresponding newborn sample, (3) a recombinant AChE protein sample, and (4) a recombinant BChE protein sample. The first two served as normalizing controls and the last two as internal controls. The iso-OMPA plates alone were incubated for 120 minutes covered in aluminum foil, and in all plates 10 µL acetylthiocholine were added to all wells immediately before measurement. Absorbance at 436 nm was read in a microplate reader (Tecan Spark, Tecan Group, Switzerland) at room temperature for 21 kinetic cycles of one minute each. The kinetic measurements were retrieved using the SparkControl Magellan 3.1 program and were further analyzed using R (46). Calculation and normalization are further available at the Supplementary methods.

### Serum RNA extraction

Short RNA was extracted from both maternal and newborn serum samples. Samples were defrosted at room temperature, 500µL aliquots were removed and centrifuged at 4°C, 3000g, for five minutes, and RNA was extracted from 200uL taken from the top of the aliquot, using the miRNeasy Serum/Plasma Advanced Kit (Qiagen, 217204) according to the manufacturer’s instructions. Concentration measurements (NanoDrop 2000, Thermo Scientific) showed average yields of 14.3 ng/µL. Quality assessment (Bioanalyzer 6000, Agilent) showed average RIN values of 1.8, reflecting the almost complete lack of rRNA in serum (47) and thus not poor quality.

### Sequencing and alignment

Of the 120 UCS samples, only those of dyads of mothers scoring either low (PSS-10 ≤ 10) or high (PSS-10 ≥ 19) PPS were considered for short RNA sequencing (Supplementary Figure 1a). The samples were divided into four groups by the mothers’ PSS-10 score and the newborn sex, and for each group the 12 samples with the highest RNA concentration in both NanoDrop and Bioanalyzer assessments were sequenced (n=48). To avoid birth types from confounding the sequencing results, we only analyzed RNA samples from non-emergency vaginal births (Supplementary Figure 1b); those 35 newborn RNA samples were divided into four groups: female and male newborns from PPS mothers (female stress, n = 6; male stress, n = 8) and female and male newborns from control mothers (female control, n = 11; male control, n = 10). A subset of 24 samples from mothers corresponding to the sequenced newborns was chosen to form matched dyads (n = 6 in all four groups), based on the highest RNA concentration in the maternal cohort.

Libraries were constructed using the D-Plex Small RNA-seq Kit from Illumina (Diagenode, C05030001) and the short RNA fraction was sequenced on the NextSeq 500 System (Illumina) at the Center for Genomic Technologies, the Hebrew University of Jerusalem. Newborn and maternal samples were sequenced on separate flow cells (NextSeq 500/550 High Output Kit v2.5 75 Cycles, Illumina, 20024906) using 1600 pg/sample in the newborn cohort and, due to lower RNA yield, 400 pg/sample in the maternal one. To compensate for the difference, sequencing depth was ∼10M reads for newborns and ∼20M for maternal samples. Subsequent quality control was re-assessed using FastQC (48) version 0.11.8, with Cutadapt (49) for removing the adaptor sequences, followed by Flexbar (50,51) for further quality screening and cleaning. Alignment to tRFs was done using MINTmap (52) to MINTbase v2.0 (53) and to microRNA using miRDeep2 (54) to miRbase 22.1 (55,56).

### Statistical analysis

#### Differential expression analysis

Differential expression analysis was performed using the DESeq2 (57) package in R with their suggested prefiltering step, disqualifying all miRs and tRFs that did not achieve rowSums(counts(dds) >= 10) >= smallestGroupSize, where smallestGroupSize=6. Normalized DESeq2 findings were deemed statistically significant for p ≤ 0.05 after FDR correction using Wald’s test. This left 565 and 174 tRFs and 69 and 49 miRs for downstream analyses in newborn and maternal samples, respectively. Analyses at the tRF family level, including length analysis, are detailed in the Supplementary Methods.

#### Identifying cholinergic targets

To identify miRs and tRFs that target genes in the cholinergic network we combined the miRDB sequence-based prediction algorithm (58,59) with a modified version of our in-house cholinergic scoring system (25–27). Briefly, we tested miRs and tRFs that were 17-28 nucleotides long who had targets with a prediction score ≥ 80, which were then matched against a list of 102 cholinergic genes (Supplementary Table 1). Those were assigned scores of one or five according to their role in the cholinergic network, with core genes (e.g. CHAT, ACHE) receiving a score of five and others (e.g. IL6, CHKB) receiving a score of one. For each miR/tRF a “cholinergic score” was calculated by summing the scores of their gene targets, thus identifying as CholinomiRs/CholinotRFs miRs or tRFs targeting (a) at least one core cholinergic genes and/or (b) at least five peripheral cholinergic genes.

#### Classification and visualization

SVM kernel classification was employed using the Caret R package (60) with a “leave-one-out” cross validation method, and the MLeval R package (61) served to calculate receiver operating characteristic (ROC) curve, area under the curve (AUC), accuracy, precision, F1 and recall values (Supplementary Table 7). P-values and subsequent false discovery rate (FDR) were calculated based on 10,000 permutations for five of the top marker groups (Supplementary Table 8).

## RESULTS

### Newborns of PPS-mothers show elevated serum AChE levels

To find if mothers’ PPS affects their cholinergic signals and influences their newborns’ reaction to birth stress, we measured serum acetylcholine hydrolyzing cholinesterase (ChE) activities in both mothers and newborns of the FELICITy cohort (40–43). We classified mothers according to their third trimester perceived stress assessment, using the PSS-10 questionnaire (7), combined with their PDQ values (44) which were predictably highly correlated (Supplementary Figure 1c). Other cohort demographic or medical features did not differ significantly between the stress and control groups (Figure 1b, Supplementary Figure 1d). Testing serum measurements of ChE activities in mothers’ serum showed similar ChE activities between the stress and control groups, possibly due to the sampling timing – upon entry to the delivery room (Figure 2d). However, newborns serum was sampled after-birth, and indeed, newborns of stressed mothers showed higher serum acetylcholinesterase (AChE) activities than those of control ones, with male newborns showing generally higher AChE activity compared to female newborns (Figure 2d; Mann–Whitney U test: Female P-value = 0.035, male P-value = 0.0048). Although not statistically significant, butyrylcholinesterase (BChE) activity in newborns showed the same trend (Figure 2d), together suggesting altered serum acetylcholine levels in stressed compared to control newborns.

**Figure 1.**
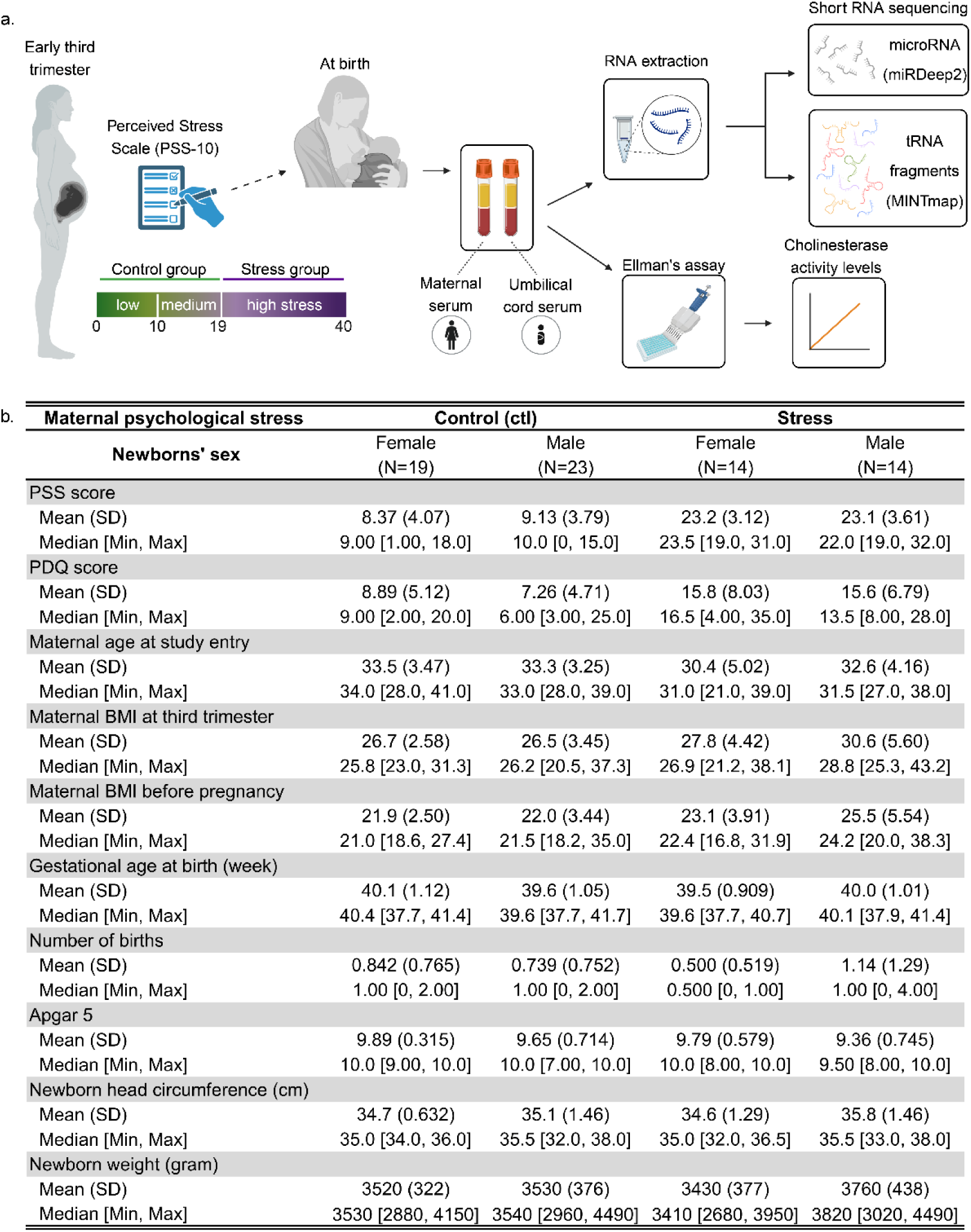
*Study layout and participant characteristics.* (a) Mothers recruited early in the third trimester were assigned to psychosocial stress and control groups based on the Cohen Perceived Stress Scale 10 (PSS-10). Maternal blood samples (n=128) were collected upon entry into the delivery room and umbilical cord blood samples (n=120) were collected postpartum. Serum was extracted and served for ChE activity measurements and short RNA sequencing. Only mother-newborn dyads who had normal vaginal deliveries were used for subsequent analyses (Supplementary figure 1). (b) The four groups reflect combinations of maternal PPS scores and newborn sex. Maternal and newborn parameters pertinent to the study are presented with both mean and median. Created with BioRender.

**Figure 2.**
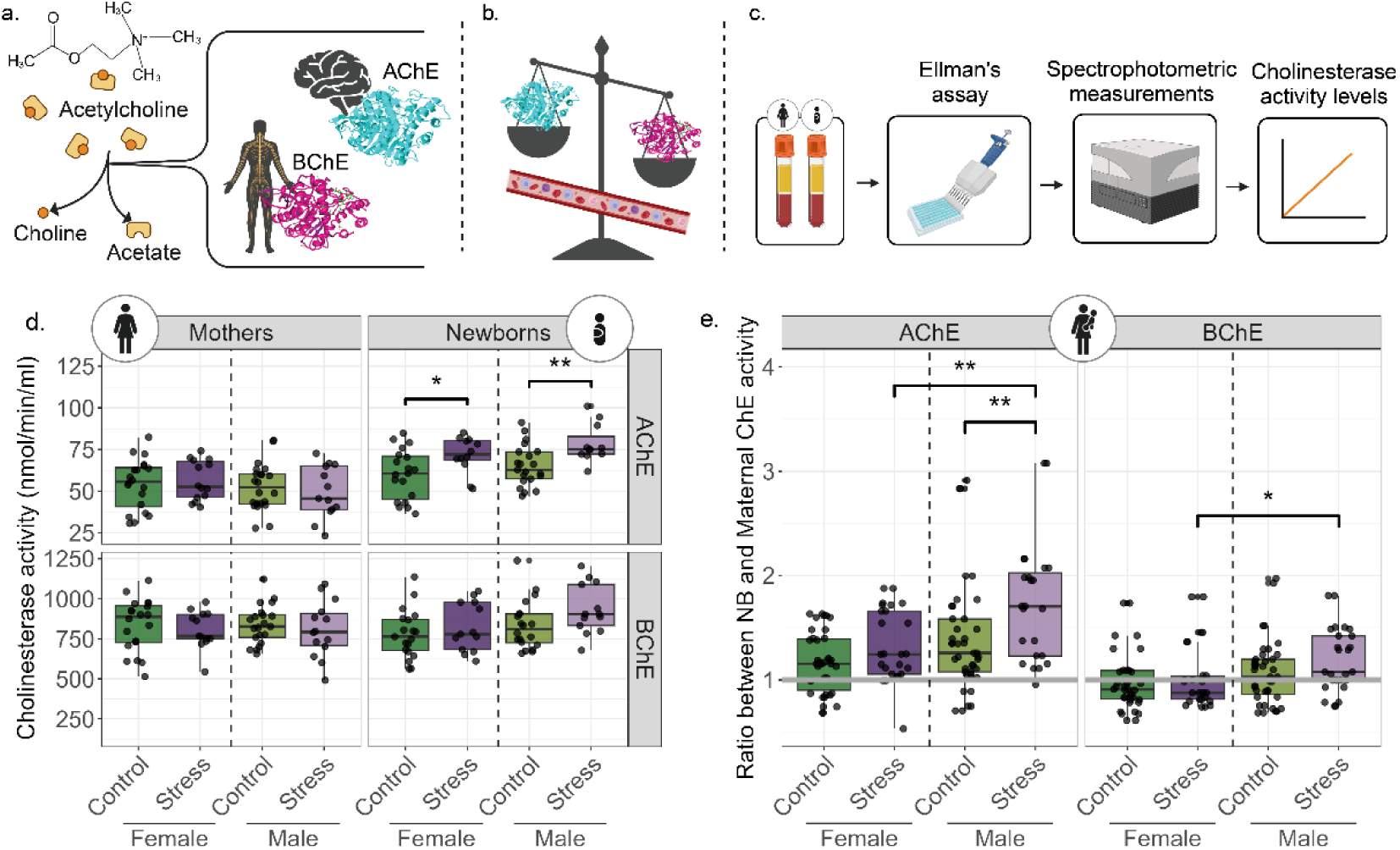
*Stressed newborns present altered ChE activities, reflecting cholinergic system involvement*. (a) The cholinesterases AChE and BChE hydrolyze acetylcholine into acetate and choline. AChE is found predominantly in the brain and BChE in the periphery. (b) Balance between the two ChEs in the blood reflects different diseases and disorders. (c) Pipeline used to assess activities of ChEs in our cohort. (d) Boxplots comparing AChE and BChE activities in the serum of mothers and newborns (AChE: mothers, n=69; newborns, n=63. BChE: mothers, n=68; newborns, n=67). (e) Ratios between newborn and mother dyads for AChE (n=62) and BChE (n=64). The grey line indicates a ratio of one (equal ChE activities in mothers and newborns). P-values < 0.05, Mann–Whitney U test. IQR method to remove outliers was used in both d and e. Created with BioRender.

Notably, serum AChE activities in all newborn groups but female controls were higher compared to their mothers (Figure 2e; Wilcoxon signed rank test, FDR ≤ 0.05; Supplementary Table 2), even when combining stress and control groups and testing all female and all male newborns compared to their mothers (female FDR = 2.9E-03, male FDR = 2.8E-04). Moreover, AChE activity was higher in stressed compared to control males (Mann–Whitney U test, P-value = 0.0072) and in stressed females compared to stressed males (P-value = 0.0087). BChE ratio presented a similar trend in the male stress group alone, showing higher BChE activity compared to their mothers (Figure 2e; P-value = 0.041; Supplementary Table 2). Hence, our findings identified higher AChE activity as an indicator of stronger response of the cholinergic system to birth stress, specifically in male newborns of PPS-exposed mothers.

### Serum miRs present small changes in newborns and mothers

Altered miRs and tRFs levels could suppress cholinesterase activities, therefore a decline in targeting miR/tRF can explain the observed elevations in AChE and BChE levels. Therefore, we compared serum miRs profiles between the newborns’ stress and control groups (Figure 3a-c). When combining male and female data, a stress-related decline in hsa-miR-301a-3p emerged (Figure 3b), predicted to control autoimmune cell differentiation through regulating STAT3 in multiple sclerosis patients (62). A parallel trend, albeit not significant, emerged in males and females studied separately (Figure 3a, c). Male newborns of PPS mothers further showed elevated hsa-miR-7704 levels (Figure 3a), which was also elevated in schizophrenia patients and may predict altered synapse activities (63). Interestingly, previous pursuit of this cohort identified one hypo-methylated CpG, cg05306225 in newborns saliva (41) that was annotated to the schizophrenia-related CSMD1 gene (64,65). Two additional miRs, the hematopoietic regulating hsa-miR-223-3p (66,67) and hsa-miR-142-5p (68,69) showed declined levels in PPS mothers of female newborns alone (Supplementary Figure 2c). Moreover, 30 of the detected miRs were expressed in both mothers and newborns, including hsa-miR-223-3p and hsa-miR-142-5p; however, hsa-miR-7704 and hsa-miR-301a-3p were unique to the newborns’ samples.

**Figure 3.**
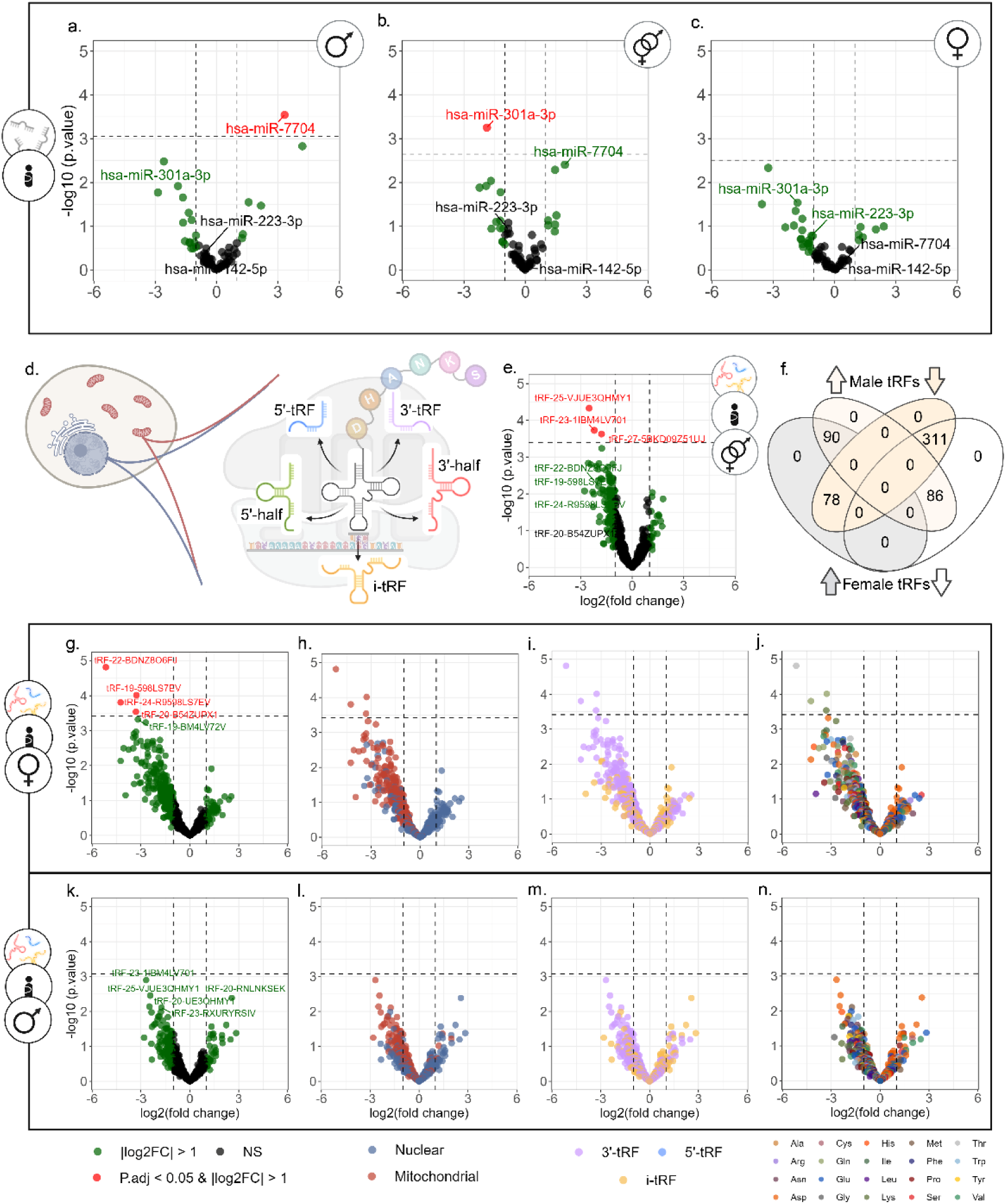
*tRF patterns differentiate stressed and control newborns, more than miRs.* (a-c) Volcano plots of miR DE analysis in three contrasts: (a) stress vs. control male newborns (n=8 vs. n=10), (b) stressed vs. control male and female together (n=14 vs. n=21), and (c) in stress vs. control females (n=6 vs. n=11). (d) tRNA genes are transcribed from two genome origins, nuclear (Nuc) and mitochondrial (MT), and encode specific amino acids. The mature tRNA is cleaved in specific locations, producing the five subtypes of tRFs: 5’-tRF, 5-half, i-tRF, 3’-half, and 3’-tRF. (e, g, k) Volcano plots of DE tRFs in (e) stressed vs. control male and female newborns together (n=14 vs. n=21), in (g) stress vs. control females (n=6 vs. n=11), and in (k) stress vs. control males (n=8 vs. n=10). (f) Venn diagram of overlapping tRFs with increasing and decreasing levels in the male and female comparisons. Volcano plots with tRFs colored according to genomic origin (h, l), cleavage type (i, m), (j, n) and tRNA coded amino acid j, n). Created with BioRender.

All four differentially expressed (DE) miRs were previously connected to the immune system and to regulating cell division in various cancers (70–77), with the two found in both UCS and maternal serum previously connected to pregnancies. hsa-miR-223-3p is highly expressed in plasma exosomes of 30 and 38 weeks normotensive pregnant women and was suggested to be one of the controllers of the fetal–maternal immune balance (78). Also, serum hsa-miR-142-5p levels were higher in mothers whose fetuses suffered from fetal congenital diaphragmatic hernia at week 26 compared to controls, and hsa-miR-142-5p levels showed a general decline between 38-40 weeks and 24 hours after birth, and was suggested to be a fetal miR that has crossed to the maternal circulation (79). These results point again to the sex-specific effect reflected by short noncoding RNAs, evident in the mothers of male newborns.

### tRF families distinguish between stressed and control newborns, with a stronger effect in females

Unlike the small PPS-related miRs differences in serum, tRFs showed more pronounced patterns, starting with three DE tRFs separating stress and control newborn groups (Figure 3e). Comparing the stress-by-sex groups revealed four DE tRFs separating female stressed newborns from controls (Figure 3g), which are intriguingly distinct from the three tRFs identified when males and females were combined. All seven DE tRFs decreased in stressed newborns. Comparing male newborns alone failed to identify any DE tRFs (Figure 3k).

Due to their shared genomic origin, parental tRNA coded amino acid, and cleavage type, diverse tRFs often present sequence similarity (Figure 3d) (20), in contrast to miRs, and may hence share similar functions. Since tRFs are simultaneously produced from tRNAs by distinct nucleases (18,19), we further hypothesized that bulks of similar tRFs may be produced and function together, creating “tRF storms” (27). If so, traditional DE analysis, which regards each tRF as an independent transcript, might fail to identify trends that would characterize whole “tRF families” whose joint biological impact may be considerable. To this end, we added family information to the tRF analysis, which identified tRF families that separate stressed and control male and female newborns (Figure 3h-j, l-n). This revealed a profound decline in the levels of mitochondrial (MT) genome-originated tRFs in stressed female newborns, with 98.9% of the tRFs showing negative fold-change (Figure 3h; exact binomial test FDR = 8.21E-74). Male newborns showed a similar, albeit less pronounced trend, with 90% of MT tRFs presenting negative fold-change (Figure 3l; FDR = 9.06E-44). Declined levels were also apparent in both sexes in 3’-tRFs (Figure 3i, m; 73.6% in females, FDR = 3.38E-18; 74.5% in males, FDR = 1.73E-19), and in i-tRFs (65.5% in females, FDR = 7.66E-06; 59.6% in males, FDR = 0.0064). tRF families grouped by their coded amino acids showed varying trends too (Figure 3j, n; Supplementary Table 2).

In contrast, the maternal cohort revealed no individual DE tRFs in any of those comparisons (Supplementary Figure 2d, h, l). Yet, tRF family patterns in stressed mothers differed from those of the newborns, revealing a decline in nuclear genome-originated (Nuc) tRFs (Supplementary Figure 2e; 71.1%, exact binomial test FDR = 2.1E-03) and i-tRFs (Supplementary Figure 2f; 80.9%, FDR = 4.5E-04), especially in mothers of male newborns (Supplementary Figure 2m-n; Nuc: 64.5%, FDR = 4.6E-02; i-tRFs: 76.6%, FDR = 2.1E-03). Further, the maternal but not newborns cohort showed globally altered levels of tRF-halves (Supplementary Figure 2f, j, n), with 3’-halves significantly elevated in all stressed mothers (87%, FDR = 2.2E-03), and in stressed mothers of male newborns separately (78.3%, FDR = 3.8E-02). Importantly, only 19 out of 45 tRF families in newborns and 30 in mothers were found in both cohorts. Moreover, the tRNA genes from which those tRFs were derived (according to the alignment by MINTmap) did not fully match the most prevalent tRNA genes, as shown by GtRNAdb (80,81) for Nuclear-originated tRNA genes and by mitotRNAdb (82) for MT-originated tRNA genes (Supplementary Figure 3a). Furthermore, tRF families with significantly modified trends did not necessarily coincide with the expected tRF families by MINTmap (Supplementary Figure 3b). This phenomenon has been seen previously (23), although to the best of our knowledge was not studied directly, and adds to the notion that tRFs production is not random but rather tRF families tend to be produced together.

### tRF families present distinct expression patterns in newborns and maternal serum

To further explore the working hypothesis that specific tRF families function as groups based on their genome origin, parental tRNA coded amino acid and cleavage type, we tested whether those tRF families show shared stress-related changes in their expression levels. Indeed, exact binomial test (FDR ≤ 0.05) identified 24 tRF families with significant DE values (Figure 4b, Supplementary Table 3). Declined levels of MT-originated Gly-i-tRFs in stressed newborns (100% in stressed newborns and in stressed female alone, FDR = 3.5E-16 for both; 96.6% in stressed male newborns, FDR = 6.1E-13) revealed the strongest signal. The nuclear genome-originated Asp-i-tRFs were strongly elevated as well (83% in stressed newborns, FDR = 0.00009; 76.6% in stressed female, FDR = 0.0025; 76.6% in stressed male, FDR = 0.009). tRF families failed to present significant changes in mothers (Figure 4b, Supplementary Table 4), however, comparing all stressed mothers to control mothers and mothers of male newborns alone revealed a trend of declined expression of Nuc-Asp-i-tRFs (92.9%, FDR = 0.055 in both comparisons). Moreover, comparing the tRF families’ profiles between mothers and newborns revealed some cases with opposite trends of change (Figure 4b), indicating non-random tRF family patterns, which called for seeking possible stress-related interaction between mothers and their newborns.

**Figure 4.**
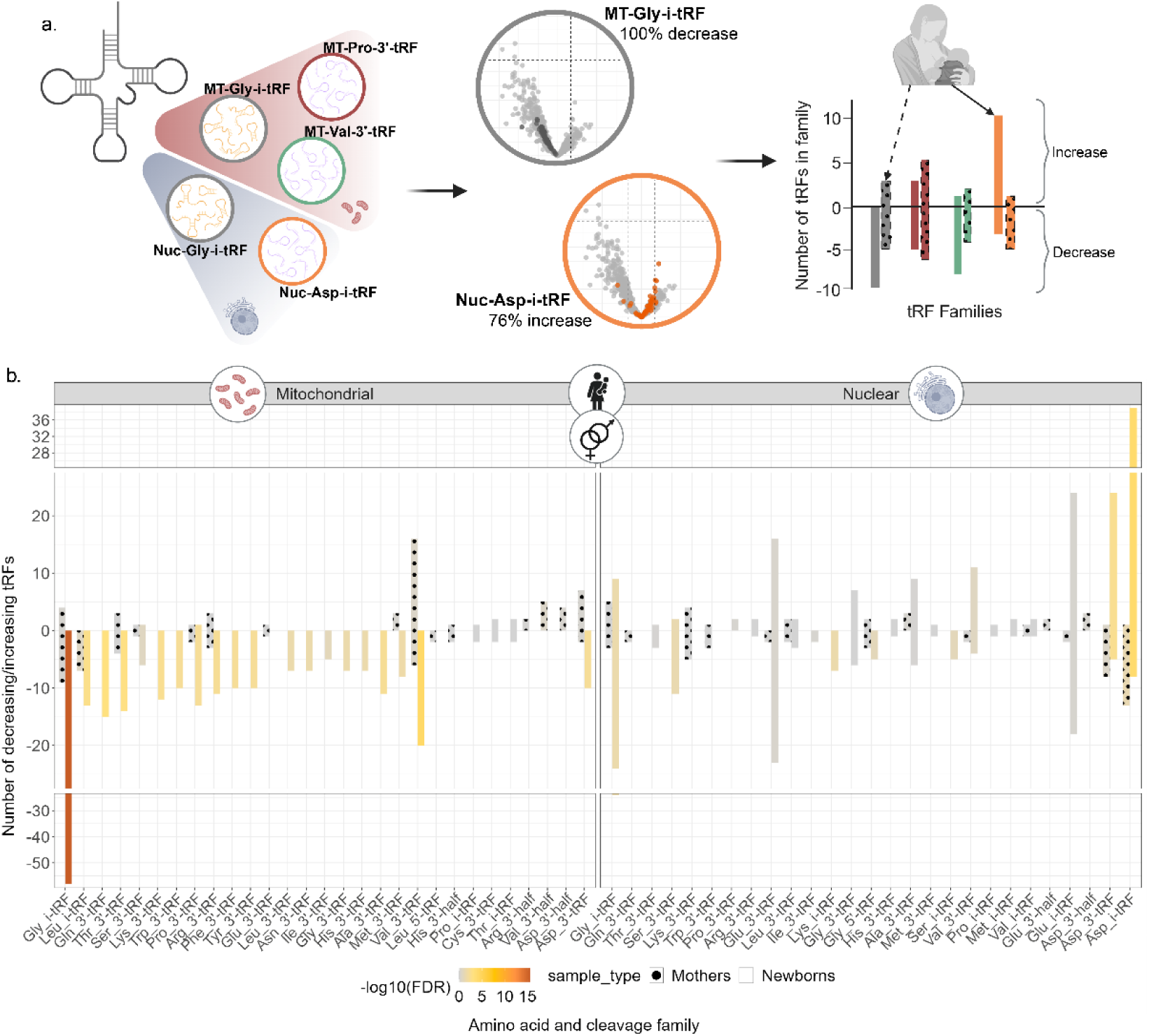
*tRF families differ significantly between stressed and control groups, with some inverse effects between mothers and newborns.* (a) tRF families were defined based on their genome origin, cleavage type, and parental tRNA coded amino acid. Only families with one or more members were included in the analysis. The Exact Binomial test (FDR ≤ 0.05) was used to quantify the tendency for decreased expression levels in the stress groups. (b) Bar plot presenting direction of change in each tRF family. Bar size reflects the number of tRFs in the identified family, with positive values indicating increased levels and negative values indicating decreased levels in the stressed groups. Bar color indicates the significance level, empty bars depict newborns and patterned bars depict mothers. The plot shows the results of males and females together. Sex-separated results are available in Supplementary Tables 3 and 4. Created with BioRender.

### tRF families show distinct stress-related length distributions

Dicer, Angiogenin, Drosha and other nucleases cleave mature tRNAs to yield the five tRF subtypes (i-tRF, 3’-tRF, etc.), producing various lengths under each cleavage category (20). To explore tRFs length distributions in our datasets, we calculated the mean expression levels of all same length tRFs in each newborn cohort (Supplementary Figure 4a). Both male and female control groups presented non-evenly distributed expression patterns, peaking around the lengths of 23-24nt, with smaller peaks around 18-19nt and 31-32nt. The highest peak in all groups emerged at the length of 36nt, but was somewhat attenuated in the female stress group, which in general showed more even distribution of tRF lengths. Intriguingly, the highest peak in the control groups overlapped known miR lengths, 16-28nt according to miRbase, (55) (Supplementary Figure 4c). In mothers, we noticed far smaller differences in length distributions between stress and sex groups (Supplementary Figure 4b), suggesting that newborns’ tRFs, more than their mothers, tend to overlap miR lengths, possibly pointing at a shared functional role of targeting and suppressing the expression of mRNAs carrying a complementary sequence (23,83). Taken together, these findings indicate that mothers’ perceived stress may lead to excessive reaction of their newborns that is reflected in declined tRFs levels.

Next, we tested whether newborn tRFs of particular families tend to appear in specific lengths and are influenced by the mothers’ PPS. For this purpose, we used the Kruskal-Wallis test on each tRFs family across the four stress-by-sex groups (Supplementary Figure 4d). In newborns, numerous tRF families showed significant preference both towards a single direction of expression change and a tendency for specific length distributions, but the maternal cohort showed no significant length differences between the groups. Notable was the MT-Gln-3’-tRF family, which showed elevated tRF lengths in both control groups compared to the stress groups in newborns, especially in females (Supplementary Figure 4e, f). Supporting previous reports of stress specificity in tRFs production (20,22,27), these findings imply that tRNA cleavage into tRFs is context-specific, presenting stress and sex group differences already at birth.

### CholinotRFs and CholinomiRs show specific cholinergic predicted targets

Since tRFs may target genes similarly to miRs and given the cholinergic stress-response network (31,32), we sought the cholinergic gene targets of the identified tRFs and miRs, combining our in-house scoring system (Figure 5a) (25–27) with the miRDB target prediction algorithm (22, 23). This identified 13 maternal and 16 newborns “CholinomiRs”, showing enrichment compared to those listed in miRbase 22.1v (Fisher’s exact test, maternal and newborn P-values of 0.007 and 0.015, respectively). In addition, we identified 13 maternal and 38 newborns “CholinotRFs”, which were enriched in MT vs. Nuc tRFs in newborns (Figure 5b; Fisher’s exact test, P-value ≤ 0.00001). These values support previous predictions in Parkinson’s and Alzheimer’s disease (25,26), deepening the connection of MT-originated tRFs with the cholinergic system under stress and neurodegeneration.

**Figure 5.**
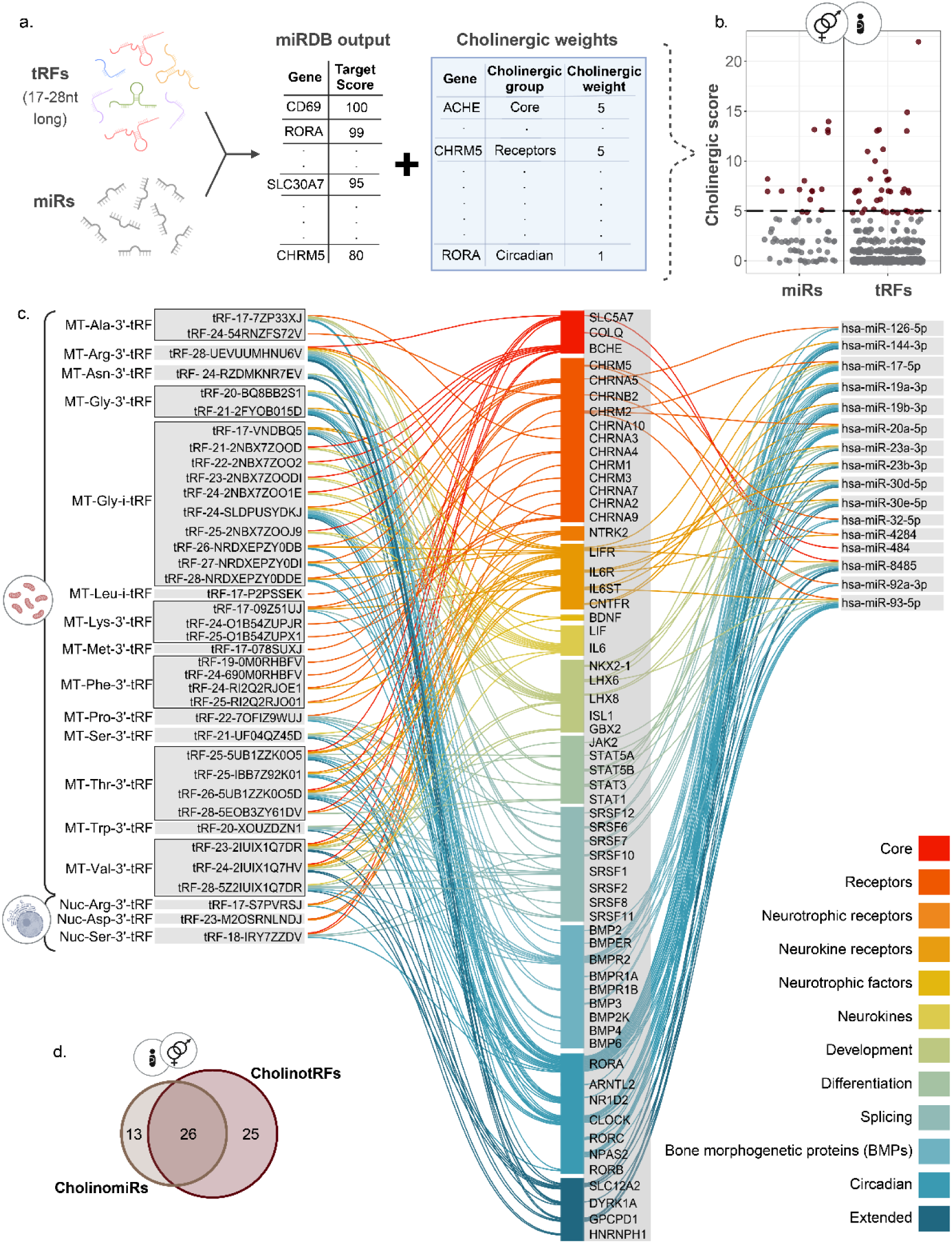
*CholinotRFs and CholinomiRs are predicted to target specific genes in the cholinergic system.* (a) Combining the miRDB sequence-based prediction algorithm with our in-house cholinergic scoring system revealed 16 CholinomiRs and 38 CholinotRFs expressed in newborns. (b) Dot plot showing cholinergic score distribution across UCS miRs and tRFs in both male and female newborns. (c) Network plot presenting predicted targets of the newborn CholinomiRs (right-hand list), and CholinotRFs which are grouped according to their tRF families (left-hand list). tRF families with more than one cholinergic member are framed by black lines. The central list of 51 cholinergic genes that are predicted targets of the CholinomiRs and CholinotRFs is colored according to the class within the cholinergic pathway. (d) Venn plot presenting the number of cholinergic gene targets shared between the newborn CholinomiRs and CholinotRFs. Created with BioRender.

Seeking overlaps between the predicted cholinergic gene targets of CholinomiRs and CholinotRFs in newborns (Figure 5c, d; Supplementary Tables 6, 7) revealed key cholinergic genes encoding enzymes such as BChE and cytokines like IL6 as targeted by CholinotRFs alone. In contrast, the circadian class of cholinergic genes and most cholinergic receptor genes were co-predicted for targeting by both CholinomiRs and CholinotRFs. However, Spearman correlations between BChE levels in newborns or mothers and its targeting tRFs did not reveal any significant correlations (Supplementary Table 5), although a correlation was found between the MT-Gly-i-tRF family and maternal BMI (Supplementary Results). Furthermore, the identified CholinotRFs belonged to 14 MT and three nuclear tRF families. Of those, only seven MT families had more than one CholinotRF member. Specifically, MT-Thr-3’-tRF family members targeted genes across the entire cholinergic landscape, whereas MT-Phe-3’-tRF family members showed specificity toward only three of the cholinergic gene classes. Likewise, the 14 newborn MT CholinotRF families presented specific length distributions, strengthening the notion of stress-driven tRF production.

### CholinotRF classify female newborns by their mothers’ PPS

To test whether short noncoding RNA levels may classify newborns into stress and control groups, we applied SVM Kernel classification with “leave-one-out” cross-validation to different sets of tRFs and miRs, testing the male and female groups together and apart, and seeking the best group of markers for this task (Supplementary Table 8). Predictably, females presented the best classification, based on the levels of all of the seven UCS DE tRFs and the four female-specific UCS DE tRFs (Fig 6a), with area under the ROC curve of 100% (AUC; 10,000 permutations, FDR ≤ 0.0001; Supplementary Table 9) and 98%, respectively (FDR = 0.008). Remarkably, the next best classification in female newborns was achieved using the 38 CholinotRFs, yielding AUC of 95% (FDR = 0.008), while CholinomiRs reached an AUC of 88% (FDR = 0.051). Male newborns, however, yielded very low classification scores, all under chance levels (Figure 6d). When comparing males and females together, testing stress vs. control groups without accounting for sex (Figure 6b) achieved classification results that were in between those achieved for male and female separately, with the best classification reached using all of the UCS DE tRFs (AUC = 78%, FDR = 0.029). Considering sex as well (Figure 6c) achieved much better classification, with the lead marker groups still being the UCS DE tRFs (AUC: Female DE tRFs = 94%, FDR = 0.008; all DE tRFs = 93%, FDR = 0.008). Combining our results with the published Fetal stress index (FSI) from the FELICITy cohort, a measurement of fetus heart rate reactivity measured non-invasively during the third trimester (43), elevated some of the classification results, mostly in male newborns (Supplementary Results, Supplementary Figure 5a-c, Supplementary Table 8, 9), further strengthening the perceived connection of cholinergic tRFs to stress and its sex-specific effects.

**Figure 6.**
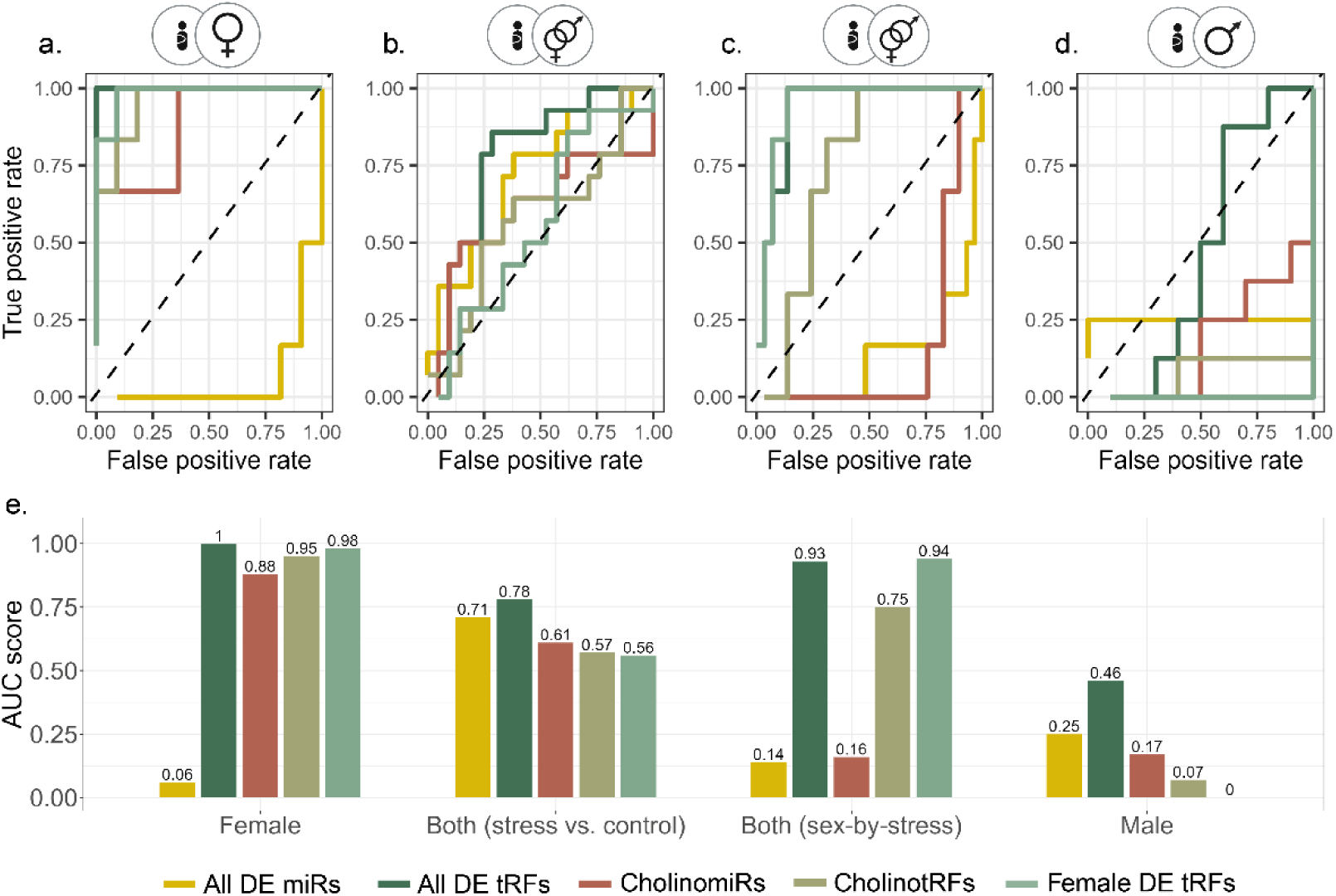
*CholinotRFs classify female newborns according to maternal PPS.* (a-d) ROC curves of five marker groups (All DE tRFs, Female DE tRFs, CholinotRFs, All DE miRs, and CholinomiRs), classifying newborns to stress and control groups based on SVM Kernel algorithm with “leave one out” cross validation: (a) females (n= 6 vs. n=11), (b) males & females (stress vs. control; n= 14 vs. n=21), (c) males & females (stress-by-sex; n= 14 vs. n=21), and (d) males (n= 8 vs. n=10). (e) Bar plot of AUC values across comparisons with FDR based on 10,000 permutations (available in Supplementary table 9). Created with BioRender.

## DISCUSSION

Maternal PPS has long been known to affect the fetus (5,8,11,12), but validated biomarkers assessing its impact remained lacking. We discovered a new group of biomarkers reflecting maternal PPS impact on the newborn, using tRF expression patterns and ChE catalytic activity levels in maternal and umbilical cord serum. Specifically, we discovered higher AChE activity in the UCS of stressed newborns, with a stronger effect observed in males that indicated cholinergic involvement. This was supported by enriched CholinomiRs and CholinotRFs showing declined levels, with the latter classifying female newborns into stress and control groups with an AUC of 95%.

Importantly, assigning these tRFs to families by their genome origin, cleavage type, and coded amino acid revealed shared expression and length patterns that were specific not only to the differences between mothers and newborns, but also between sex and stress conditions, where female newborns of PPS-affected mothers showed the largest differences.

Our findings have a dual value. First and foremost, we present tRFs as novel biomarkers that differentiated male and female newborns according to maternal stress assessed at birth. Further, the stress-specific tRF patterns primarily consisted of a major decline of MT tRFs that occurred in both sexes but was more profound in female newborns. Although our cohort is unique, it is small, including only 35 sequenced samples, and due to low RNA quantities, we were not able to validate our findings by qPCR measurements in the rest of the cohort. Furthermore, we did not find publicly available datasets that were similar enough by tissue and species to validate our results, most likely due to the complexity of generating a similar cohort. However, previous animal studies support our observations: Bartho *et al*. (84) injected pregnant mice with cortisone between E12.5 and E14.5, which led to a decline in mitochondrial DNA in the placenta of female compared to male offspring; and Su *et al*. (29) found the overall percentage of MT tRFs in a murine prenatal immune activation model to be higher in the fetal brain and liver of females compared to males.

The magnitude of the MT tRF effect can also reflect the use of serum rather than whole blood or PBMCs. Serum RNA reflects the extracellular vesicles and broken blood cells content, including platelets which are rich in mitochondria (85). While serum RNA amounts are low (86), which requires more specialized library preparation kits, the enrichment in short RNAs compensates for the low yield (86,87). Further, serum is the standard blood fraction in use in hospital settings and therefore forms an available pool for RNA-seq clinical studies.

Serum is also a standard tissue for measuring cholinergic tone, defined as the balance between AChE and BChE, both of which hydrolyze acetylcholine to acetate and choline. AChE acts as the main cholinesterase of the central nervous system and BChE of the periphery (31,32). In the fetal environment, besides the erythrocyte-bound blood AChE (38), AChE is found in the placenta and in umbilical cord endothelial cells (88,89). Indeed, we found higher AChE levels in the UCS compared to the maternal serum, followed by stress-related elevation in newborns. This effect was more predominant in male compared to female newborns, compatible with reports of larger stress-related differences in male newborns (5,14,42). By hydrolyzing acetylcholine, the two ChEs can suppress its anti-inflammatory effect, activating pro-inflammatory pathways including the NF-κB pathway which contradicts the vagus-mediated acetylcholine secretion in response to an activated HPA (32–34,90). The regulation of this cascade by short noncoding RNAs (38,90) echoes our findings of the seemingly unrelated enriched CholinotRFs and CholinomiRs and the two ChEs in our data. Furthermore, transgenic excess of the AChE-targeting CholinomiR miR-132 was incompatible with fetal survival in mice (91); also, cholinesterase enzymes in male mice were directly influenced by prenatal physical and psychological stress, and hyperexcitability of the cholinergic neurons in the latero-dorsal tegmentum may cause anxiety-like behaviors and memory impairment (92). Secondly, and as important, is our novel interpretation of tRF sequencing results. Although sets of tRFs share many characteristics that warrant their assembly into groups, they are still perceived by the scientific community as separate entities. Thus, while the literature often mentions tRFs by their “family name” (MT i-tRF^HisGTG^, Glu-50tsRNA-CTC, etc.) (93,94), they are still referred to as single tRF molecules. In contradistinction, our classification into “tRF families” based on a combination of their genome origin, cleavage type and coded amino acid, has highlighted expression and length patterns that are context-specific, i.e., influenced by stress and sex. Furthermore, previous studies have identified “tRF storms”, characterized by rising levels of tRFs that replaced miRs two days post-stroke (27). Indeed, in our data we saw that these family patterns passed the boundary of single parental tRNA gene and were shared even between all of the tRFs of the same coded amino acid. This marks the importance of mass production of tRFs (22) and strengthens the concept that tRFs function as groups rather than as individuals.

To the best of our knowledge, reference to tRF groups appeared in partial ways in three previous studies. The most recent, by Akins et al. (95), examined multiple datasets seeking 5’-halfs and long 5’-tRFs in human patients and cell lines. They showed that while context-dependent, tRFs that share length and parental tRNA may also share expression levels and sequence. However, their DE analysis did not examine tRF families as groups directly, and their focus was on 5’ -tRFs and -halves of specific lengths, 27-41nt, which were almost completely absent in our data. Su et al. (29) investigated the influence of prenatal maternal immune activation in mice on tRFs in several tissues. Immune activation was performed at E11.5 and E12.5 and within a few hours after the second activation they thoroughly investigated the tRF landscape, including an in-depth characterization of the length distribution according to cleavage type by both sequencing and northern blot analyses of tRFs grouped by their parental tRNAs. However, using STAR alignment (96) rather than the standard annotations of tRFs, rendered their analysis less translatable by the standard tools of the field. The third study, by Isakova et al. (97), aligned tRFs with a combination of STAR and Unitas (98) too, as one part of a complex atlas of short noncoding RNAs in different murine tissues. Grouping tRFs based on their genome origin, cleavage type, and coded amino acid codon, showed tissue-specific expression patterns, however, their only statistic for assessing tRFs as groups was the sum of expression, overlooking the variability within said groups. Further, i-tRFs were absent from their analysis.

Finally, all three papers retained groupings based on single amino acid codons, while we have shown that similar expression patterns were shared even within the wider boundaries of a single amino acid coded by several codons. Our introduction of the binomial test allowed for the quantitative examination of how tRF families function as groups, by testing shared expression patterns, length distributions and sequence similarities. Nevertheless, we wish to stress out the importance of previous reports establishing specific functions of single tRFs in diverse diseases and conditions (23,25–28,30), and urge future researchers to take these points into account and seek both single and familial tRF functions.

In conclusion, we have shown the involvement of PPS in affecting the cholinergic system at diverse levels, by modulating newborns ChE activities, CholinomiRs and CholinotRFs. Our study demonstrated the importance of tRF families as sex-specific biomarkers that reflect the effect of maternal PPS on newborns, as early as the birth itself. We presented a new analysis approach that considers the group function of tRFs as families, as determined by their genome origin, cleavage type, and coded amino acid, rather than as single tRFs. We believe that these findings play an important role in the early diagnosis of stress effects in newborns and encourage further research into the landscape of tRFs in prenatal and other stress-related conditions.

## ACKNOWLEDGEMENTS

The authors wish to thank all participants and all the medical staff at the labor and delivery of the Department of Obstetrics and Gynecology (Klinikum rechts der Isar, Technical University of Munich, Germany) for their valuable contribution to the study. The research leading to these results received funding from the Israel Science Foundation, ISF (835/23; to H.S.), the European Research Council (Advanced Award 321501, to H.S.), and from a joint research support to H.S. and L.C. from The Hebrew University. We further acknowledge the Ken Stein foundation’s support. L.C. is the Snyder Granadar chair in Genetics, H.S. is the Slesinger chair in Molecular Neuroscience. This work was further supported by a Hans Fischer Senior Fellowship from IAS-TUM (Institute for Advanced Study - Technical University of Munich, Munich, Germany) awarded to M.C.A., as well as funds from the Department of Obstetrics and Gynecology, Klinikum rechts der Isar, Technical University of Munich, Germany to S.M.L., and CIHR to M.G.F. (grant number 123489).

## ETHICS APPROVAL

The study protocol is in strict accordance with the Committee of Ethical Principles for Medical Research from TUM and has the approval of the ‘Ethikkommission der Fakultat fur Medizin der Technische Universitat Munchen’ (registration number 151/16S). ClinicalTrials.gov registration number is NCT03389178. Written informed consent was received from participants prior to inclusion in the study.

## DATA AVAILABILITY

The data underlying this article will be uploaded to GEO.

