## Supplementary material for "Maternal prenatal stress induces sex-dependent changes in tRNA fragment families and cholinergic pathways in newborns": Supplemetary Information

#### Supplementary Information

### SUPPLEMENTARY METHODS

#### Cholinesterase activity calculation

Average values of "mean OD/minute" units were converted to “nmol substrate hydrolyzed per minute per ml” units using the Beer-Lambert law (A= εlc; molar absorptivity constant ε 2-nitro-5-thiobenzoate= 13,600), multiplied by the serum dilution factor (1:20). Normalization between plates was based on the maternal and newborn control samples described above and was performed separately for maternal and newborn samples. Values were log-transformed to minimize the effect of outliers on the mean. The normalization factor of plate $i$, denoted $f_i$, is computed by

$$f_i = \frac{\sum_{j=1}^{n} \log\left(control_j\right)}{n} - \log(control_i),$$

where $n$ is the number of plates. Let $s_{i,k}$ be the activity of the non-control sample $k$ in plate $i$. The normalized activity, denoted $s'_{i,k}$, is computed by

$$s'_{i,k} = s_{i,k} \cdot e^{f_i}.$$

#### Assignment of tRF families

tRFs share sequence similarities due to their shared tRNA origin and production mechanism (1). In our study, we used these characteristics as grouping factors, assigning different tRFs that share genome origin (mitochondrial or nuclear), coded amino acid (as referenced by the parental tRNA gene), and cleavage type (5'-tRF, 5-half, i-tRF, 3-half, 3'-tRF) (2), to the same “tRF family". Using the multiple sequence alignment (MSA) algorithm in the msa R package (3) we sought sequence similarities of all tRF families represented in our newborns data which had more than one member (45 out of 51 families). Of those families, 43 enabled us to produce a consensus sequence that coincided with a real tRF, and our data included 42 of these “consensus tRFs” that were 16-27nt long (Supplementary Table 10, Supplementary Figure 6). It is important to note that we did not find a sequence motif that was shared between all tRF families. Furthermore, unlike the nuclear genome that contains multiple tRNA genes for each amino acid, the mitochondrial (MT) genome carries only one tRNA gene for each amino acid, apart from leucine and serine with two each, and each tRNA can encode groups of four closely related codons due to an unmodified uridine at the wobble position among other related mechanisms (4,5). As the majority of our data was comprised of those MT tRFs which are coded in the MT-DNA mostly by a single codon, it sufficed to classify tRFs according to the parental tRNA coded amino acid (Supplementary Figure 3a). Even so, we could show that multiple tRF families constructed from nuclear tRNA genes representing different codons showed similar sequences and united expression patterns (Supplementary Figure 6, Figure 4). In addition, length distribution analysis was performed as follows: mean expression levels of each tRF family under each length it spans was calculated for each participant. Next, Kruskal Wallis test was used for each of the tRF families separately, followed by FDR correction, to compare their mean expression between the stress-by-sex groups. Together, these steps enabled us to assess the differences between specific tRF families rather than individual tRFs.

### SUPPLEMENTARY RESULTS

#### MT-Gly-i-tRF family correlates with maternal BMI

Searching whether the expression of certain tRF families correlated to participant characteristics such as gestation age, newborn weight etc., revealed a strong correlation between the MT-Gly-i-tRF family and maternal BMI. 40 of the 59 members in this family showed high positive correlation between the tRF expression levels and maternal BMI during the third trimester, and 35 also correlated with the mothers’ body weight at the same time point (Supplementary Table 5). Interestingly, the interaction between Glycine (Gly) and weight has been shown previously, with circulating Gly showing an inverse correlation with metabolic disorders such as obesity, non-alcoholic fatty liver disease, and type 2 diabetes (6–8). Moreover, Gly was shown to be important in late stages of pregnancy (9), and one of its related tRFs, the nuclear 5’-tRF-GlyGCC, was proposed as a regulator of obesity-associated pathways in human breast cancer cell lines, as well as mice and pigs (10,11), altogether suggesting that these findings were not coincidental.

#### Third trimester Fetal stress index (FSI) enhance classification results in male newborns

Interestingly, combining our results with the previously published FSI from the FELICITy cohort, a measurement of the fetus heart rate reactivity measured non-invasively during the third trimester (12)(Supplementary Figure 5a) elevated some of the classification results (Supplementary Figure 5d-h, Supplementary Table 8, 9), although it was not correlated with any of the tRFs or miRs and by itself did not yield more than 48% AUC. This was especially the case in male newborns, where combination of the FSI and all four DE miRs from UCS and maternal serum achieved an AUC of 95% (FDR = 0.058; Supplementary Figure 5g), an elevation of approximately 60%. In females, CholinotRFs together with FSI had an increase of 5%, reaching an AUC of 100% (FDR = 0.008; Supplementary Figure 5d). A similar elevation was observed when separating stress and control groups of males and females together, with the combination of FSI and all DE tRFs yielding an AUC of 83% (FDR = 0.029, 5% elevation, Supplementary Figure 5e). Most of the other marker sets did not do so well with the addition of FSI, showing decrease in classification success (Supplementary Table 8, 9). Together, these findings indicate that newborns can be differentiated by their mothers' PPS already at birth, as reflected by UCS short noncoding RNAs, with the sex of the newborn playing a significant role in determining the type of markers required.

### SUPPLEMENTARY FIGURES

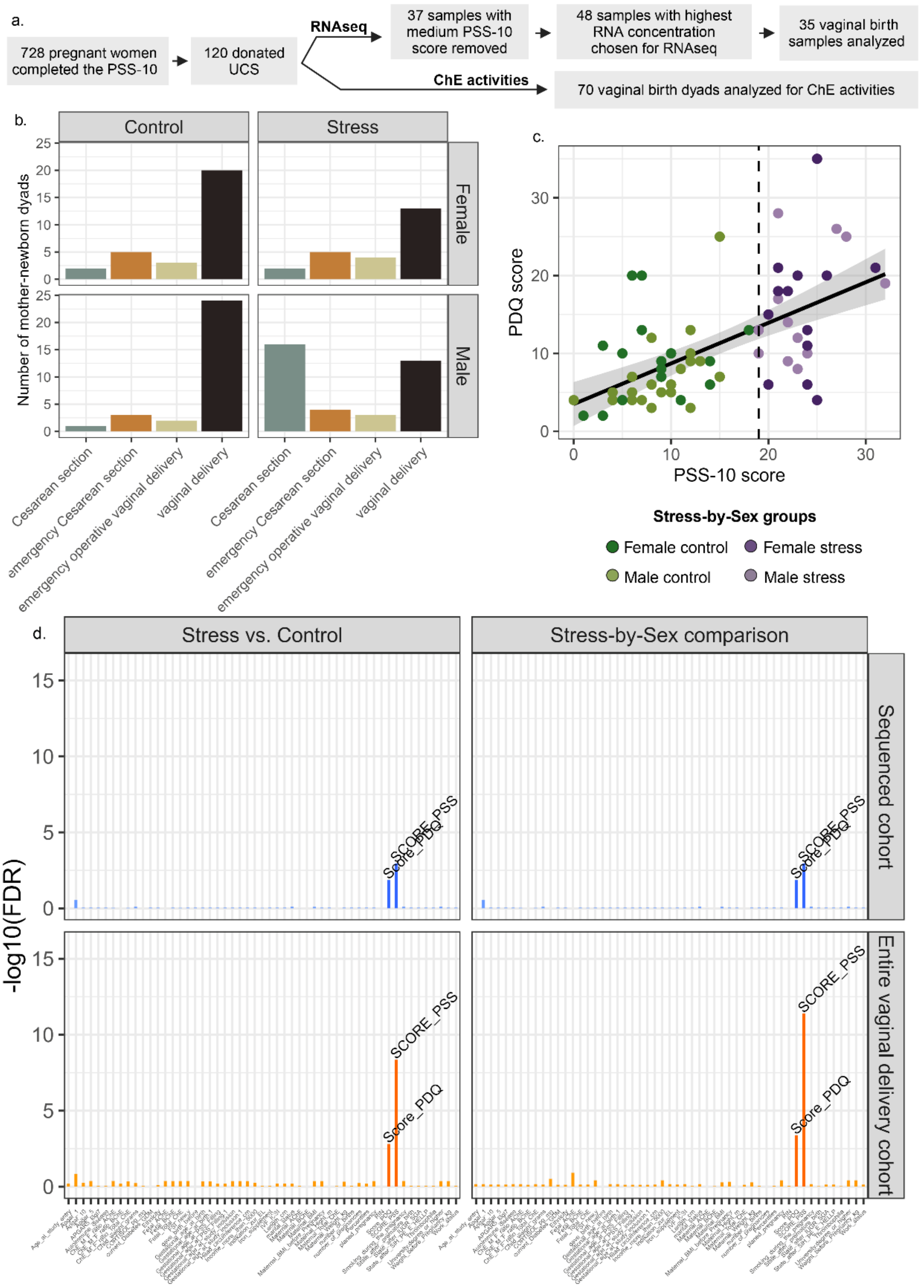

*Supplementary Figure 1* - **Data characteristics.** (a) Of 728 pregnant women filled the Cohen PSS-10 during their third trimester, 120 gave UCS at birth. Next, only samples of dyads with either low (PSS-10 ≤ 10) or high (PSS-10 ≥ 19) PPS were considered for short RNA sequencing. In each of the four groups, the 12 samples with the highest concentration in both Nanodrop and Bioanalyzer assessment were chosen for sequencing (n=48). Post-sequencing, four samples were disqualified due to extremely low/high counts and finally only samples from vaginal birth dyads were analyzed. ChEs activities were measured from all 70 vaginal birth dyads. (b) Bar plot showing the distribution of birth type in the four study groups, assigned according to the mothers' PPS by PSS-10 and the newborn sex. 58.3% of 128 mothers experienced normal vaginal delivery. (c) PSS-10 and PDQ show significant positive correlation across individuals (Pearson R = 0.6, P-value = 5.3e-08). (d) Bar plot presenting testing features across groups using Kruskal-Wallis test with FDR correction. Only the two stress-assessing questionnaires (PSS-10 and PDQ) were significantly different between the groups. Created with BioRender.

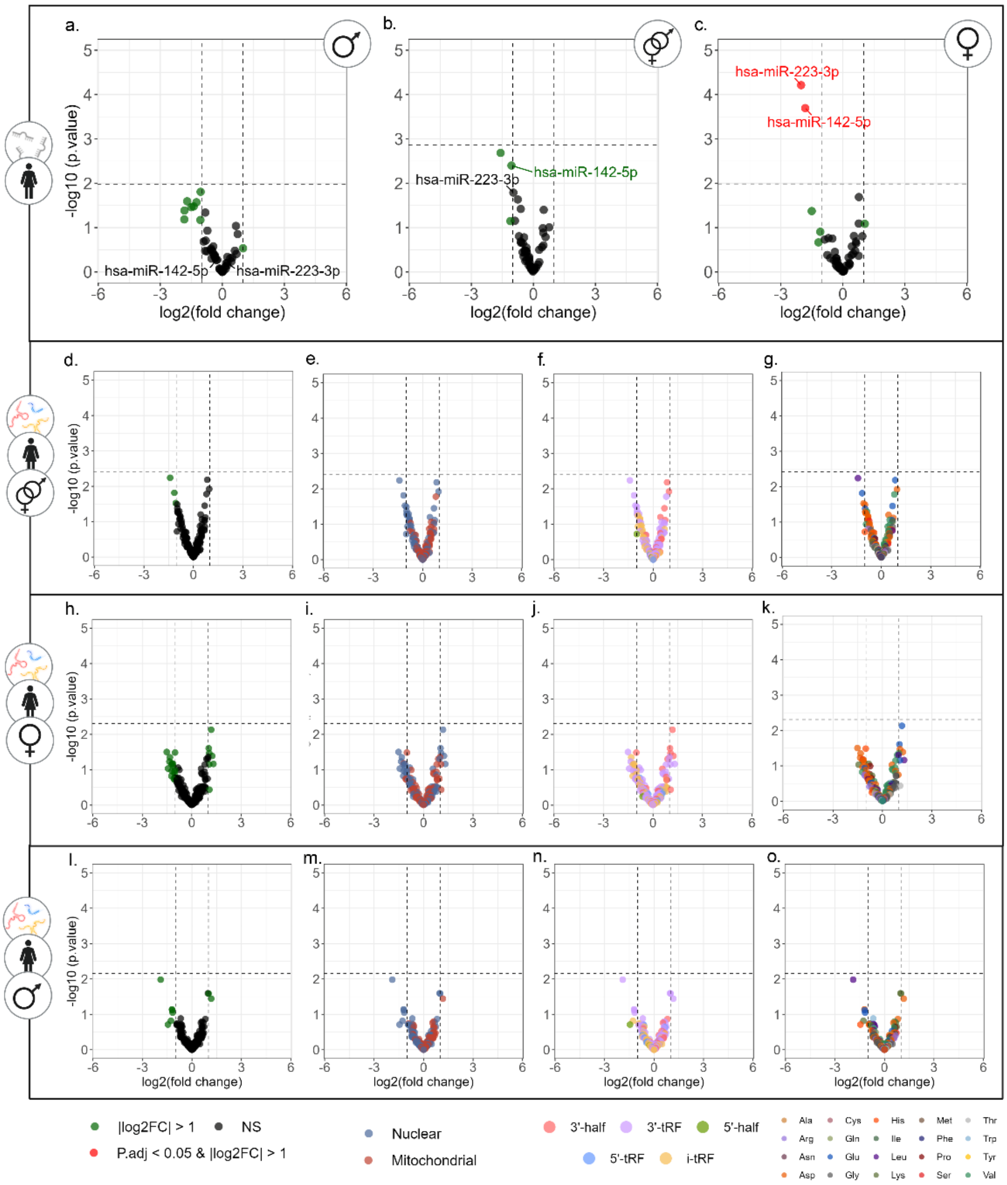

*Supplementary Figure 2* - **Maternal tRF and miR profiles do not present similar effects to newborns.** Volcano plots of DE miRs (a-c) and tRFs (d-o), in (b, d-g) PPS vs. control mothers of male and female newborns (12 vs. 12), in (c, h-k) PPS vs. controls mothers of female newborns (6 vs. 6), and in (a, l-o) PPS vs. control mothers of male newborns (6 vs. 6). tRF Volcano plots colored according to (e, i, m) genome origin, (f, j, n) tRF cleavage type, and (g, k, o) coded amino acid. Created with BioRender.

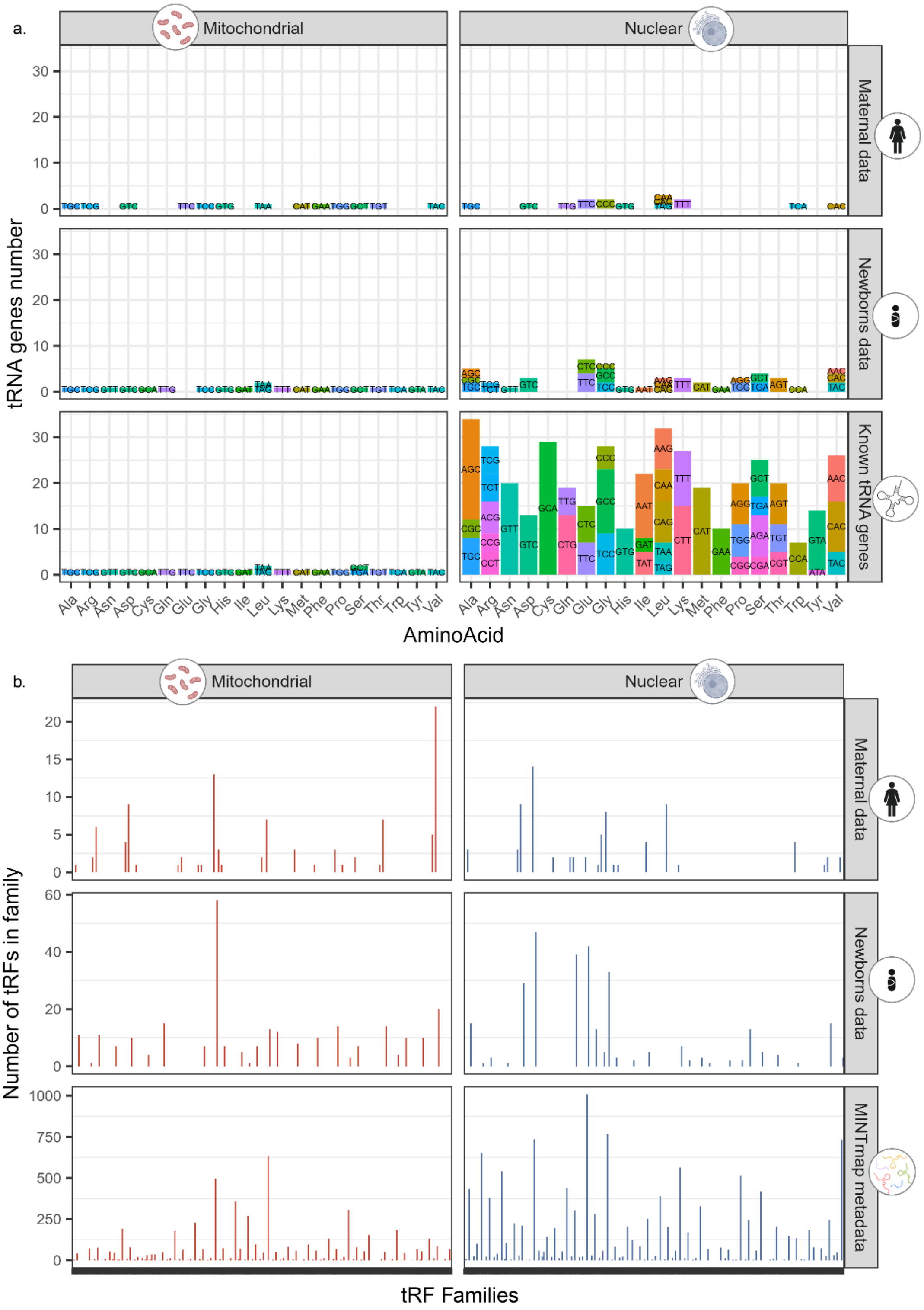

*Supplementary Figure 3* - **tRNA genes and tRF families' distribution across data sets.** (a) Bar plot presenting the distribution of tRNA genes represented in our data compared to known tRNA genes. The left Y-axis shows the number of tRNA genes, on the right the division by cohort: mothers, newborns, and known tRNA genes by GtRNAdb for Nuc tRNA genes and mitotRNAdb for MT ones. The top X axis divides tRNA genes into MT and Nuc genome origin, and the bottom to the coded amino acid. The bar color depicts tRNA codons. (b) Bar plot presenting tRF families' size, grouped based on genome origin, cleavage type, and coded amino acid. The left Y-axis shows tRF numbers in the family, and the right division to cohorts:

mothers, newborns, and all possible tRFs according to the combined metadata files by MINTmap. The top X axis shows division to MT and Nuc genome origin and the bottom to tRF families. Created with BioRender.

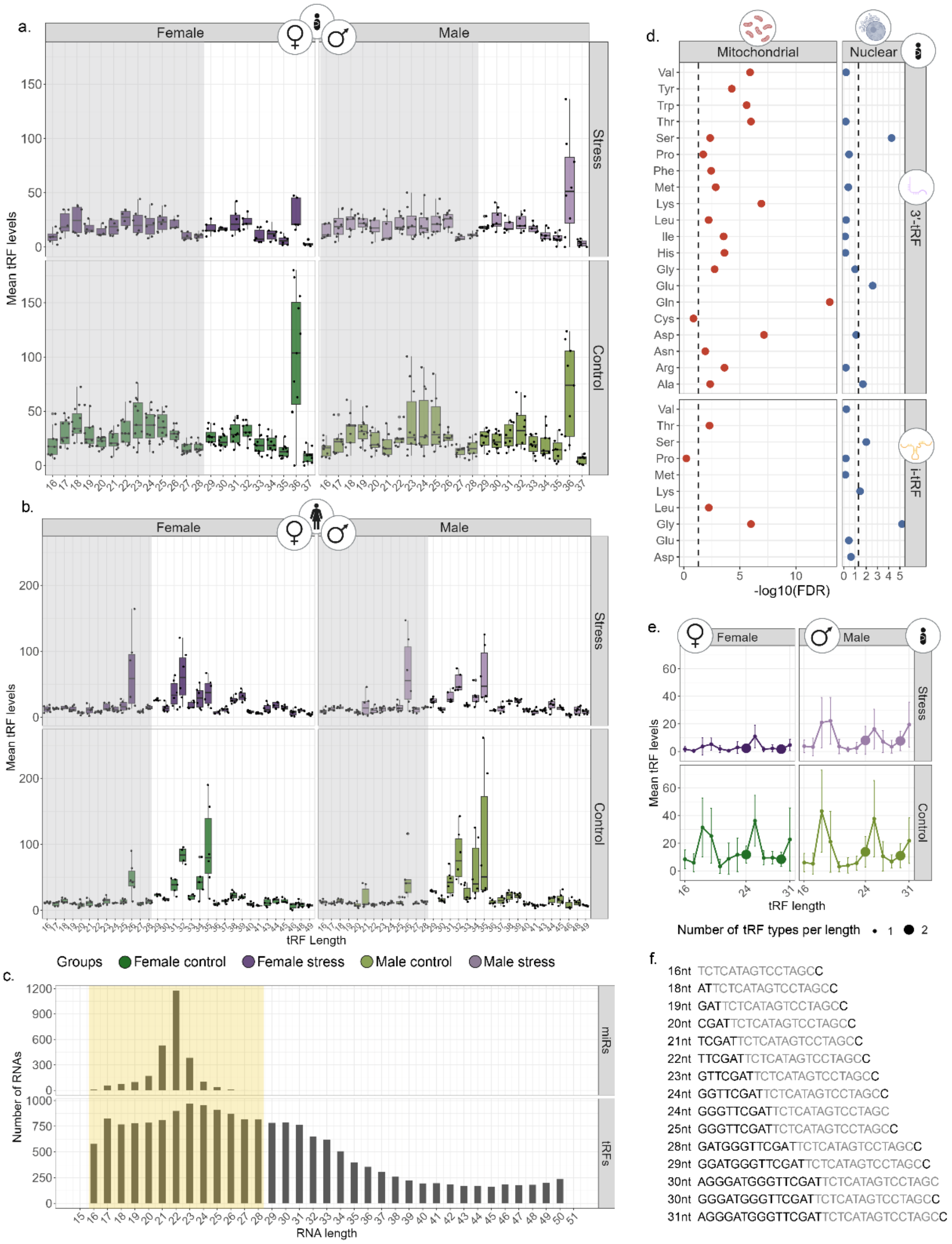

*Supplementary Figure 4* - **tRF families differ in length across newborn stress and sex groups.** (a) Boxplot showing mean expression levels of all tRFs of each length, calculated separately for the newborns of the four stress-by-sex groups (n=35), with outliers identified and excluded from the analysis by the IQR method. The black dots represent mean values for single participants. The grey areas are lengths that overlap with miR length according to miRBase (16-28nt). (b) the same for the mothers (n=24). (c) Length distribution of miRs and tRFs in known data sets. The x-axis shows lengths in nucleotides, y-axis shows the number of RNAs. miRs length distribution is taken from miRbase, and tRFs length distribution is taken from MINTbase metadata. (d) Kruskal-Wallis (KW) test of length distributions between the four groups, for each tRF family, of nuclear and mitochondrial origin. The dashed line marks the threshold of significance (FDR = 0.05). (e) Dot plot of the KW test for length distribution of the top tRF family, MT-Gln-3'-tRF, in each of the four groups. The length of tRFs in the family is plotted vs. their mean levels and the dot size reflects the number of tRFs sharing the same length in that family. (f) The sequences of all MT-Gln-3'-tRF family members in our data, from the shortest to the longest. Grey letters mark the sequence shared by all members. Created with BioRender.

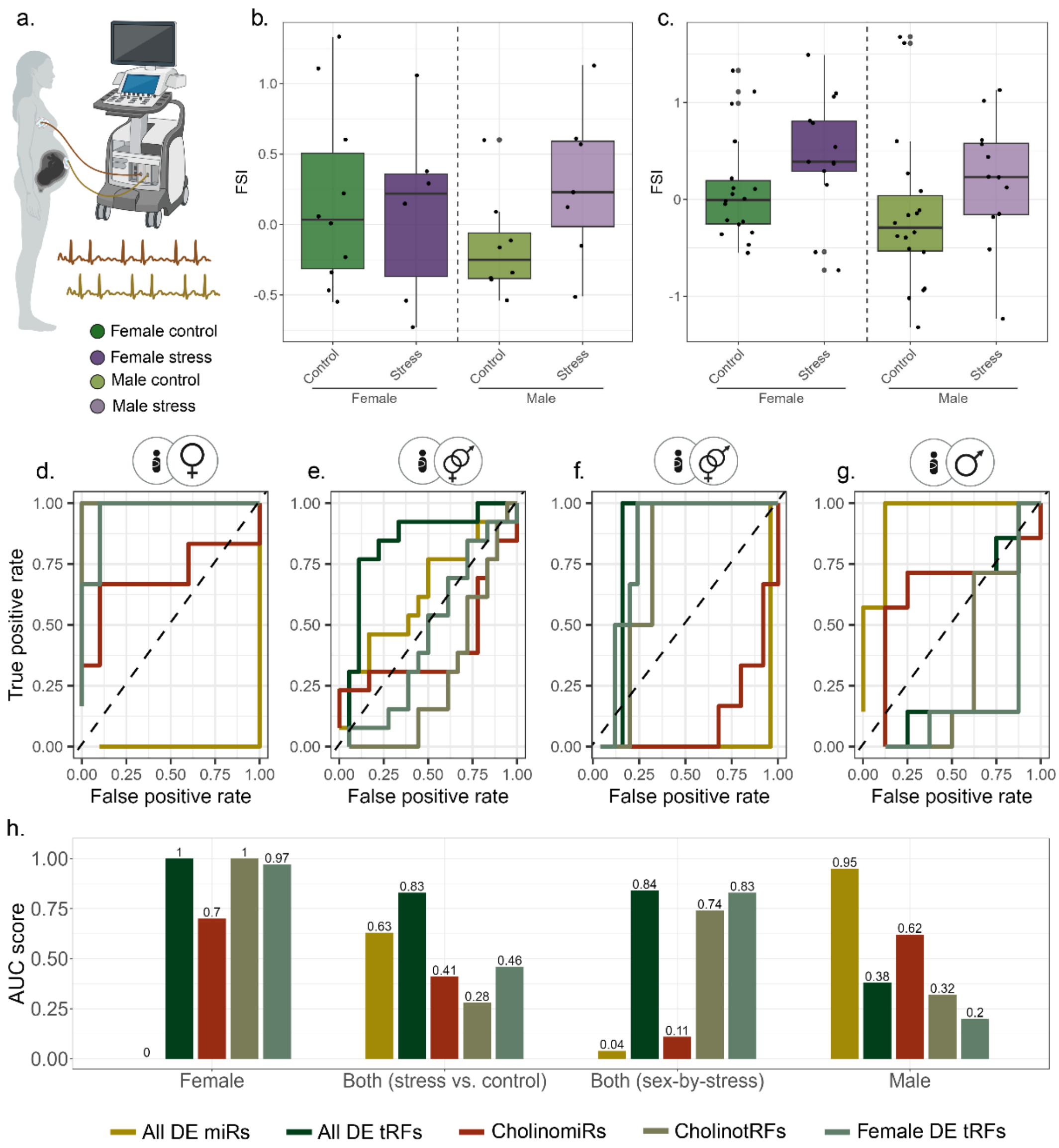

*Supplementary Figure 5* - **FSI increases the accuracy of PPS classification of some of the marker groups.** (a) FSI measurement of the fetus's heart rate reactivity, measured non-invasively during the third trimester. (b-c) Boxplots of FSI results, (b) in the sequenced cohort (n = 35 dyads), and (c) in the entire cohort of only vaginal deliveries (n = 70 dyads). None of the comparison was significant, although comparing female newborns stress and control dyads came close (u.test, P.value = 0.062). (d-g) ROC curves of five marker groups (All DE tRFs, Female DE tRFs, CholinotRFs, All DE miRs, and CholinomiRs), classifying newborns to mothers' PPS and control groups based on SVM Kernel algorithm with "leave one out" cross validation: (d) females (n= 6 vs. n=11), (e) male & female (stress vs. control; n= 14 vs. n=21), (f) male & female (stress-by-sex; n= 14 vs. n=21), and (g) males (n= 8 vs. n=10). (h) bar plot of AUC values across comparisons (FDR based on 10,000 permutations available at Supplementary Table 9). Created with BioRender.

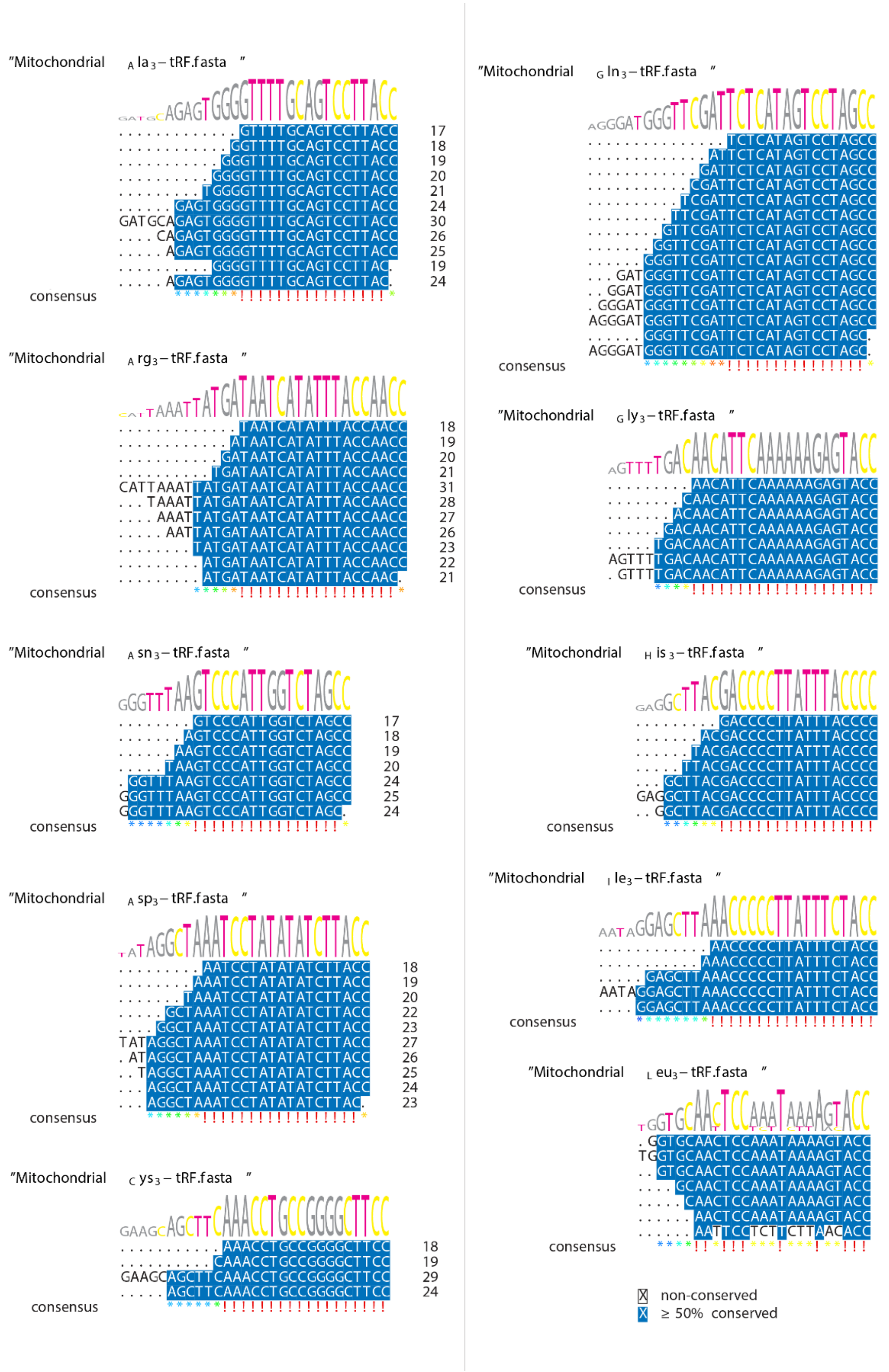

*Supplementary Figure 6* – **tRF families share sequence similarities.** Each box shows the multiple sequence alignment (MSA) of all the tRF members in our data that belong to the same tRF family. Each line shows the relevant sequence of a specific tRF in the family, with the numbers on the right side indicating the length of each tRF. White background shows non-conserved nucleotides and blue shows conservation in more than 50% of sequences. The consensus sequence is shown on top of each box. A total of 48 families are depicted over pages 10-15.

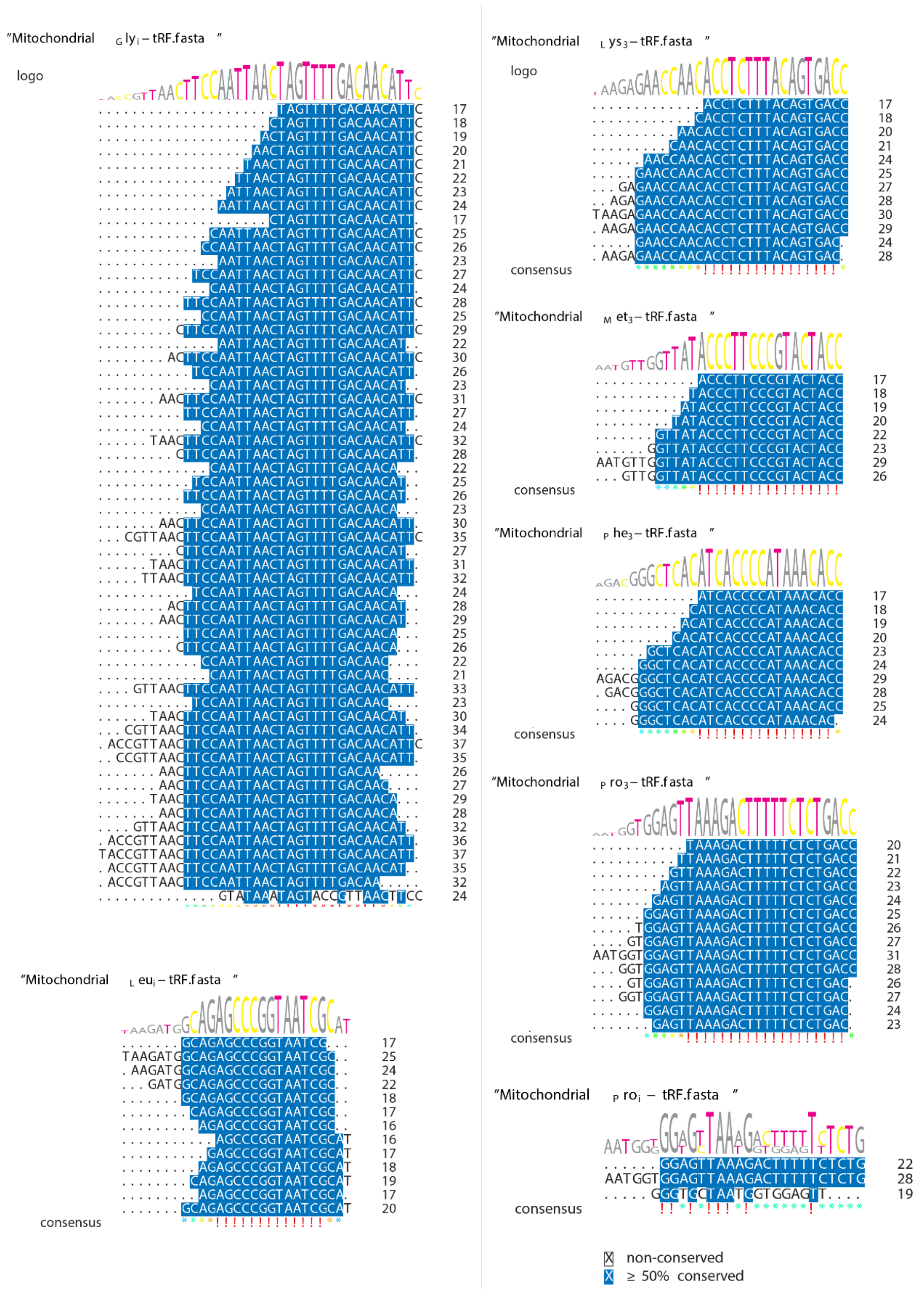

*Supplementary Figure 6 –* **continue.**

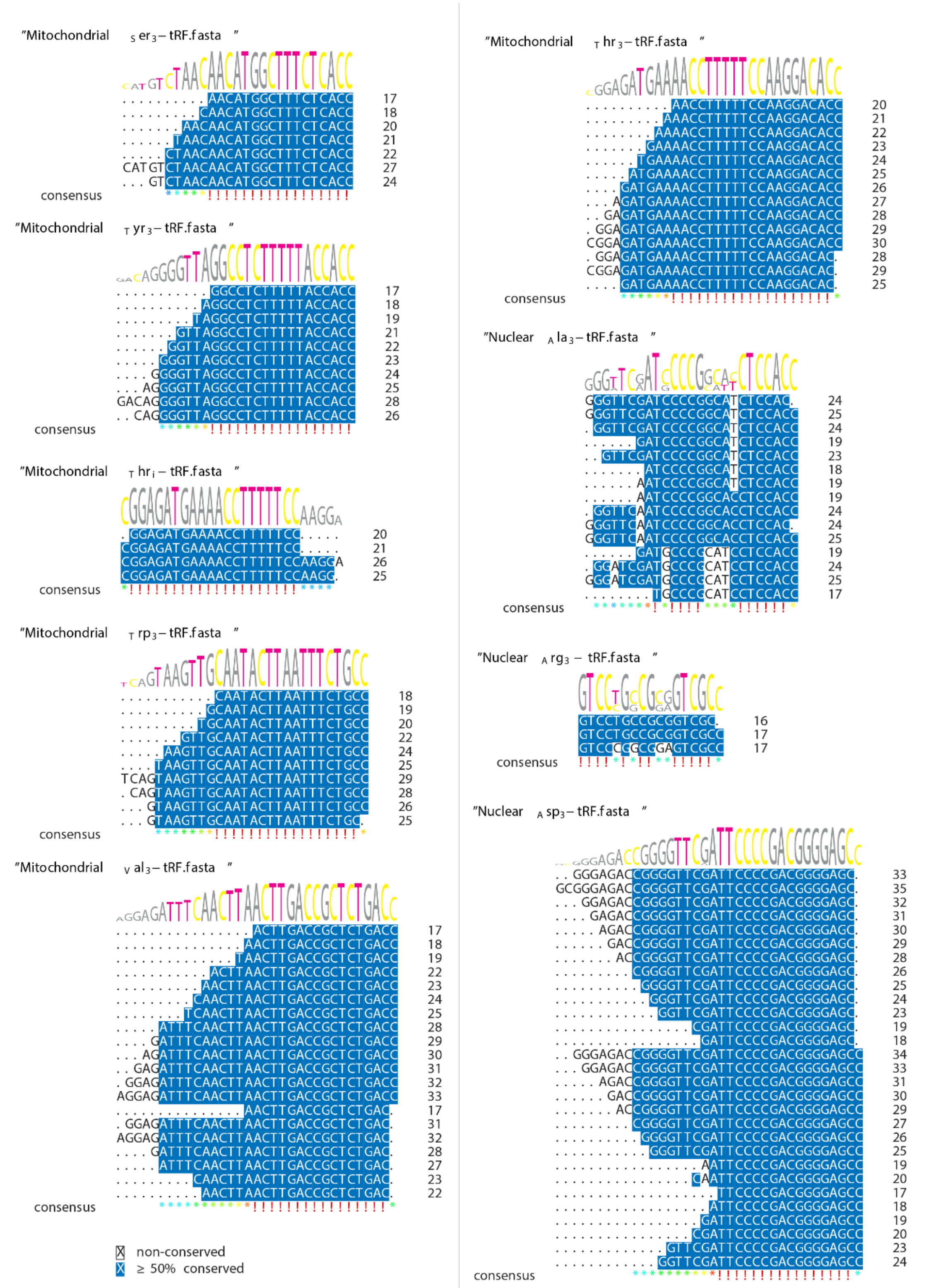

*Supplementary Figure 6* – **continue.**

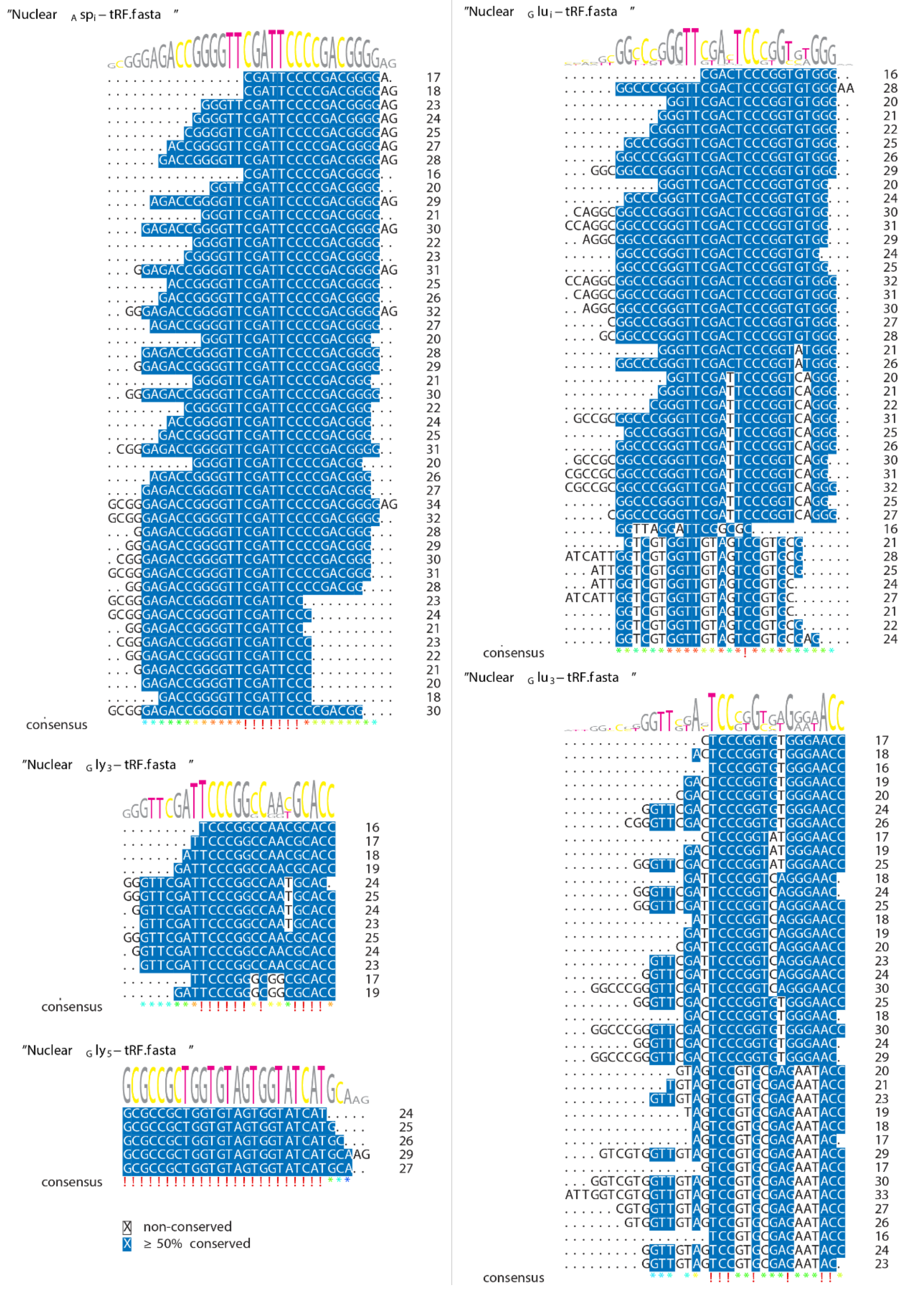

*Supplementary Figure 6 –* **continue.**

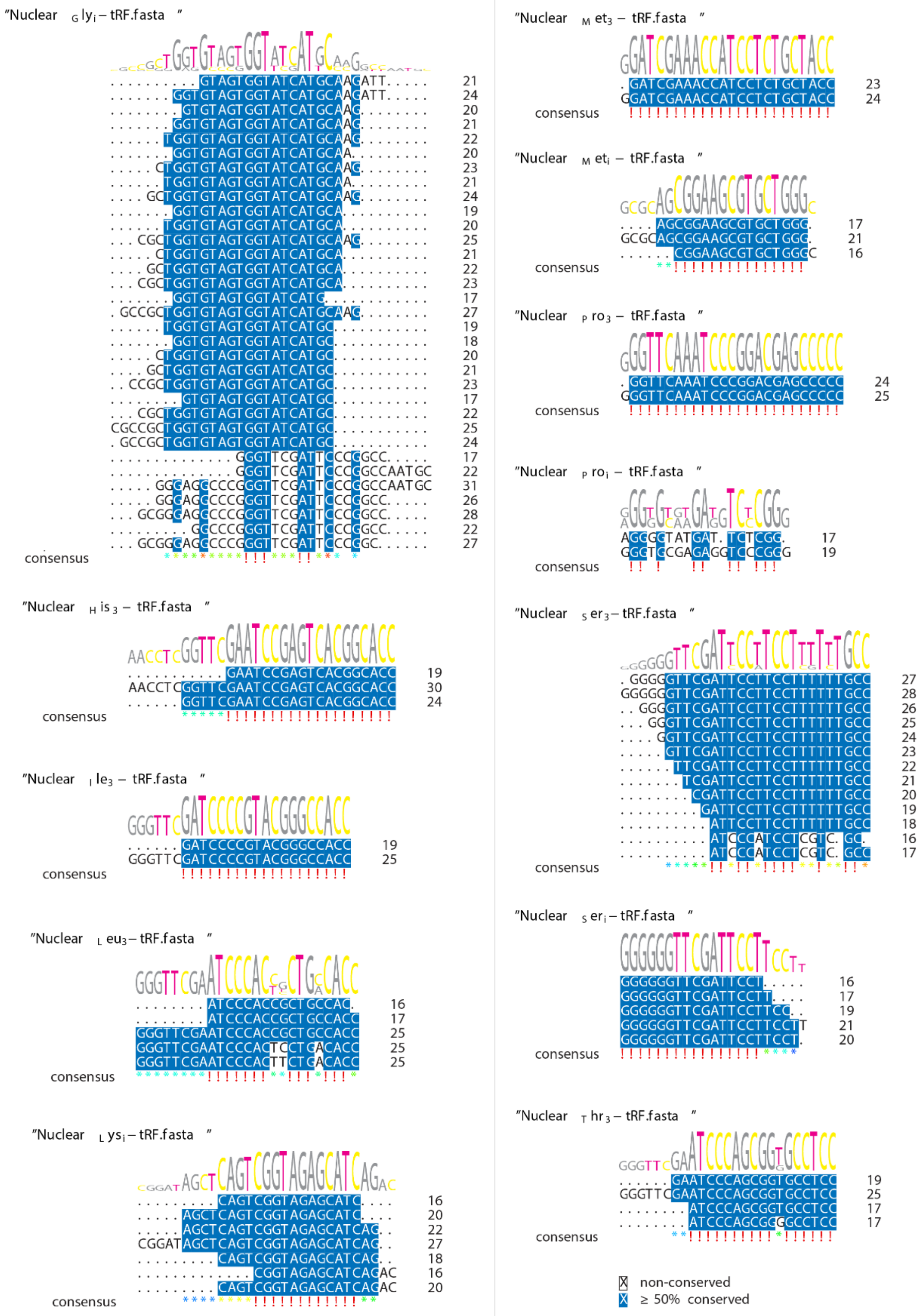

*Supplementary Figure 6* – **continue.**

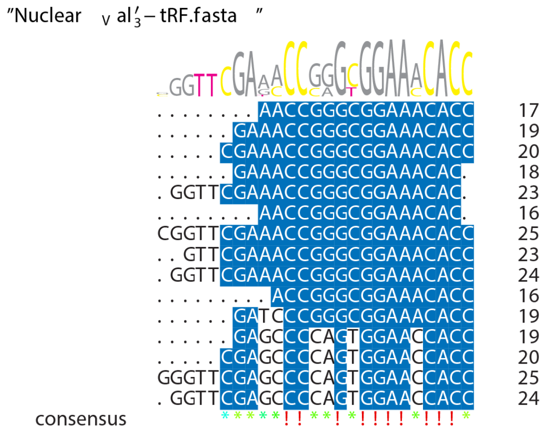

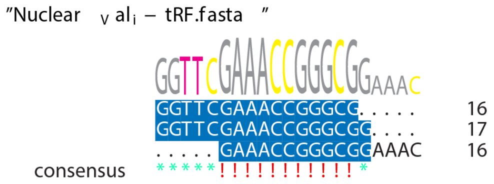

X non-conserved

X ≥ 50% conserved

*Supplementary Figure 6* – **continue.**
